# Unsupervised protein language models learn patterns of enzyme function

**DOI:** 10.64898/2026.04.23.720319

**Authors:** Matthew Penner, Michal Lihan, Hannes Bormke, Peter Nix, Hanna Moscho, Paul Dupree, Florian Hollfelder

## Abstract

While enormous amounts of sequence information have become available, assignment of sequence to a particular enzymatic function has remained elusive. Here we describe a framework that drives a general protein language model to find a target reaction without specific training, using an initial bridgehead protein. At the heart of this framework is PLM-clust, an algorithm that employs k-means on top of protein language model embeddings to convert sequence space into functional reservoirs of latent space, and samples from these clusters based on accelerated zero-shot scoring. We demonstrate PLM-clust in a recursive discovery process (with enzyme hit rates quickly rising to >90%), segmenting isofunctional reservoirs and exploring them in greater detail. This approach – exemplified for glycosyl hydrolases (a xylanase, >100-fold activity increase) and for imine reductases (IREDs, >100-fold increase in catalytic promiscuity profiles) – reliably brings about novel enzymes that are proficient at the catalytic task at hand, reaching deeply into sequence space with a majority of residues exchanged.

## Introduction

Accurate zero-shot assignment of the relative propensity of an uncharacterized enzyme to catalyse a given reaction, starting merely from information on sequence, the target transformation and reaction conditions, remains important^1^ but challenging^2^. Progress is hindered by the enormity of sequence space, the non-linear effect of mutations when combined together^3^, and the challenge of representing a chemical reaction and target conditions in a meaningful universal language^4^. This means that exploration of function in sequence space must be grounded and guided by accumulation of experimental data^5^. However, compared to *in silico* functional prediction, experimental data is slow and expensive to acquire, so the suggestion of enzymes to be tested should be kept to a minimum informative set. We propose a solution to this problem via the workflow shown in **Figure 1**.To obtain an enzyme with a target set of properties, a **p**rotein **l**anguage **m**odel (PLM) is used to embed sequences in search space nearby to a known ‘near-miss’ candidate, and these embeddings are clustered using unsupervised learning. A member from each cluster that is predicted to have high fitness for its natural function by zero-shot scoring is selected and experimentally characterized for the function(s) of interest. Now the groundwork for recursion is laid: function is projected from the labelled representative sequence to a whole cluster. Clusters with high performance for the target function are used as inputs for another round of the cycle. Each cycle therefore requires only as many experiments as there are clusters. In this way, embedding space can be traversed towards the enzyme in the dataset with optimal target function. This approach has the advantage that the target reaction conditions are imposed on PLM-clust at the experimental stage, meaning that the experimentalist has an arbitrary number of degrees of freedom in reaction condition specification. In contrast to a directed evolution campaign (Figure 1h), this approach is not hindered by fitness valleys in the fitness landscape.We sample diverse embedding spaces (Figure 1i) and aim to reach global fitness maxima otherwise might unreachable due to epistatic constraints on the protein’s evolution or accessible only via long mutational path to the optima^6^.

**Figure 1:**
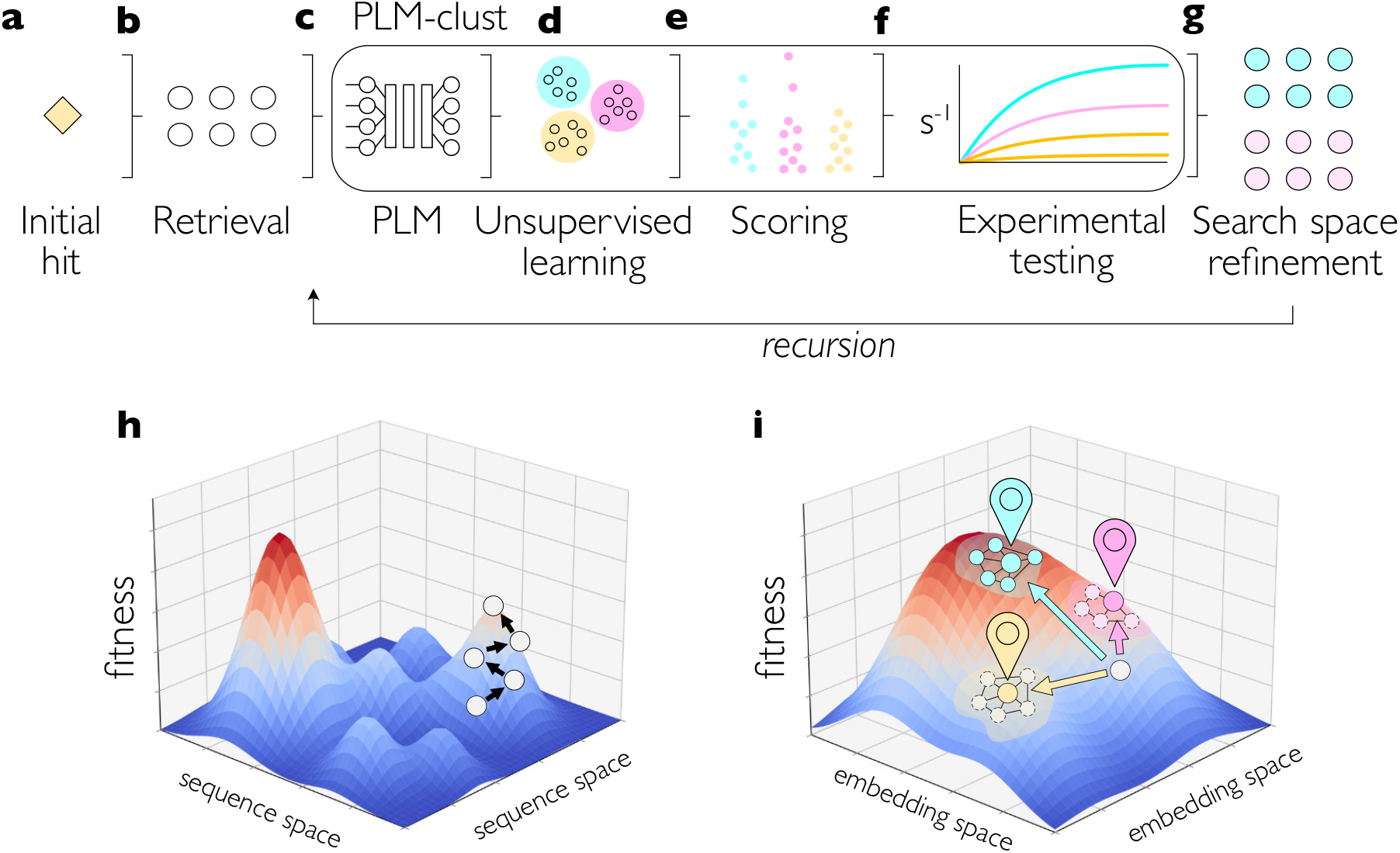
Mining embedding space using PLM-clust: providing access to novel enzymes with minimal experimental effort. (a) An initial hit, as defined by an enzyme that has been experimentally established to have some fitness for the task at hand, is selected. It is used for (b) retrieval of homologues to establish an initial search-space.These homologues are each (c) embedded using a protein language model (PLM), and these embeddings are (d) assigned clusters using an unsupervised learning technique. Members of each cluster are (e) scored using a zero-shot filter, and the top scorer from each cluster is (f) produced and experimentally characterized for the function of interest.The top performing enzymes are used to (g) identify clusters that are enriched in the function of interest, which are then used in a recursive call to the PLM-clust algorithm. Typically, approximately 10 enzymes need to be experimentally screened with high confidence (∼90%) that predictions can be produced in quantity. (h) schematic of traditional stepwise directed evolution on a narrow area in a sequence landscape taking a starting point up to a local fitness peak. (i) schematic of how PLM-clust aims to traverse an ‘embedding landscape’, starting with the same starting point, PLM-clust can sample embedding space (pins) even if the fitness peaks are separated by low-fitness valleys. Further iterations exploit the area around newly discovered functional space (blue points), eventually covering a wider area in sequence space than in (h).

This framework mirrors the generation and use of well-established Sequence Similarity Networks (SSNs)^7^. Enzyme discovery using SSNs involves generating all-versus-all comparison of the sequences of interest based on local homology and/or alignment coverage (e.g. using BLAST^8,9^), and defining a relatedness threshold, above which two sequences are considered ‘linked’ in the network (resulting in an ‘edge’ in the visual 2D representation, outlined in SI Figure 1). There are principled methods for selecting this threshold based on metrics such as closeness centrality that help an expert define a meaningful relatedness threshold^9^, but methods based only on protein alignments miss out on context critical to decoding protein function, such as residue co-dependencies, structural information, inter-protein interactions^10^, and the relevance of each residue in the protein to the protein’s function. Although SSNs are convenient tools for segmenting sequence space, and widely used as a proxy for functional similarity^7,11,12^, they are very sensitive to choice of input parameters, relying on expert knowledge on the enzyme family to construct^9^, and fail at ‘long-range’ similarity tasks not captured by a no-longer-meaningful alignment^13^. Alone, they are insufficient for predicting substrate specificity^5,14^, leading to high experimental failure rates when predictions are tested. In contrast, pre-trained protein language models (PLMs) present as a more information-rich starting point for enzyme discovery. Trained on unlabelled protein sequences and capturing positional conservation, coevolution, and structural constraints, PLMs learn an internal representation of proteins called an *embedding* that embodies these properties across the primary sequence^15^.These models are trained on sequence context, meaning that amino acids are interpreted not as independent entities, as in an MSA, but all together in the context of the protein in which they are found^13^. PLMs can even bypass the need for human-defined substitution matrices to represent each amino acid, by learning an encoding for each amino acid during the training process, and using it as a ‘first guess’ before passing the whole protein through the model and moulding the embeddings to their sequence context^13^. In preliminary investigations of how embeddings correspond to enzyme function, Shaw et al.^16^ have shown that language model embedding similarity can be more informative than sequence similarity on the ability of protein homologues to complement each other between organisms. PLMs can also be used for zero-shot prediction of protein ‘fitness’,^17^ and while the exact definition of fitness learned by a protein language model is as yet unclear, it likely relates to how well the patterns found in a protein correspond to patterns seen across the evolutionary training data. This is tangible enough to predict experimental mutational profiles across a wide range of proteins and tasks^18^.

One of the problems inherent to using protein language models to compare proteins is that embedding size is dependent on protein length^19^. Comparing embeddings of different sizes requires a decision about how to handle this mismatch, a problem conventionally addressed with alignment-based methods using the concept of ‘gaps’^8^. Detlefsen et al.^20^ have previously overcome this problem by transforming the primary embedding output of a PLM to a length-independent embedding to represent all sequences in a length-invariant manner, while Shaw et al.^16^ have used Wasserstein embeddings, as defined for PLMs by NaderiAlizadeh & Singh^21^. To this end we use mean-pooled embedding (MPEs), where a sequence is transformed to a single ‘average amino acid’ embedding vector. Compared to other methods that yield length-independent results (such as concatenation and bottlenecking)^20^ information may be lost, but this approach does not require representation fine-tuning on the query set, and parameterization is not required between different enzyme targets^20^, allowing comparison at both short and long range in sequence space. For analogous reasons, mean-pooling has been applied successfully to language models in other fields, such as natural language processing.^22^

Here, we have investigated how the pretrained PLM ESM2-650M can be applied to the task of catalytic activity mining (Figure 1), and how when combined with unsupervised learning, it can guide an experimentalist towards stable, active enzymes with target function matching or exceeding a known enzyme, used as a starting point for such a campaign. We validate our approach using two examples: first, the discovery of biocatalysts, and secondly, for mining promiscuous activities in a model enzyme family. We go on to show that this framework can be generalized to other PLMs, connecting future advances in self-supervised machine learning models to application in enzyme discovery.

## Results

### MPE generation followed by analysis of local neighbourhoods show that neighbours in embedding space are also neighbours in catalytic function

To test the ability of MPEs to account for alignment ‘gaps’^8,23,24^ in diverse datasets, we probed whether embeddings of enzymes with similar catalytic function would occupy similar regions of MPE-space, even with low sequence similarity. As an example, we took a sequence- and structure-diverse subset of the expert-curated CAZy (carbohydrate-active enzyme) database^25^ and extracted hydrolase and transferase enzymes at random.We then applied ESM2 to generate MPEs for each enzyme.Visualization of the results with a t-SNE captures the local relationships in the data.This striking split between two functional CAZyme classes, hydrolases and transferases that occupy different regions of MPE space (Figure 2B), suggests that functional relatedness is preserved: hydrolases are nearest-neighbours to other hydrolases in 96.9% (13851 of 14294 cases). As glycosyl hydrolases and glycosyl transferases occupy a diverse array of folds, this indicates that properties beyond fold, structure, and sequence similarity are captured by this method of embedding protein sequences.A control experiment indicates that these similarities are not captured by purely sequence-based methods, as shown by the sequence-similarity network of the same data (Figure S1). Here, similar sequences are indeed clustered but reflect fold similarities detected by high sequence homology. Across folds, sequence identities <30%, are insufficient to detect fold similarity^26^ and catalytic similarity across folds is not captured. Indeed, it is not possible to generate a meaningful SSN for such a diverse dataset, with over 60% unlinked members (Figure S1), in comparison to 96.9% of enzymes sharing a nearest neighbour with another member with their hydrolase or transferase annotation using PLM-clust.

**Figure 2:**
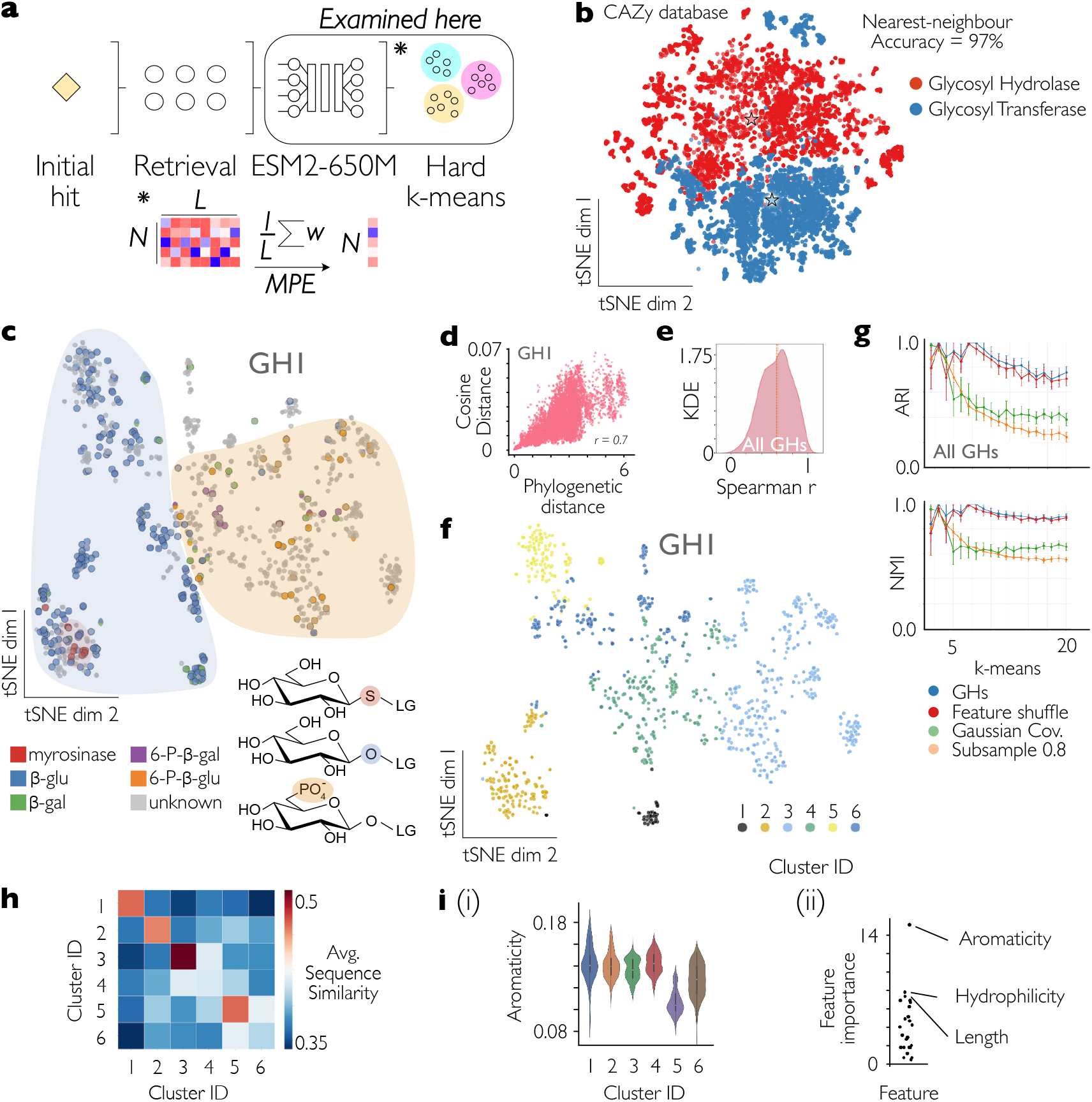
The embedding space of ESM2 reports on both structure and function in a well-annotated database of enzymes with diverse functions. **(a)** The head of the PLM-clust workflow. A set of target sequences is processed by the ESM2 protein language model, resulting in one full-length LxN (L = protein length, N = model dimensions) embedding per sequence; the full-length embeddings are averaged across the length of the protein and mean-pooled embeddings are obtained; the mean-pooled embeddings are clustered using hard k-means clustering, here with three means displayed **(b)** A representative subset of the carbohydrate-active enzyme database (CAZy) visualized using mean-pooled ESM2 embeddings and a t-SNE. Sequences are coloured according to their general catalytic function – glycosyl hydrolases (GH) compared to glycosyl transferases (GT). Sequences with multiple annotated catalytic functions have been removed for clarity, as these often represent multi-domain enzymes. Stars indicate cluster centroids. **(c)** The family GH1 represented in ESM2 embedding space – sequences with similar annotated catalytic function cluster in similar regions of high-dimensional space, as represented with a t-SNE and coloured by EC class, as extracted from the CAZy database. Representative substrates are shown, with their distinguishing features highlighted. LG = leaving group. **(d)** Phylogenetic distance derived from a multiple sequence alignment (MSA) of the GH1 family (N = 10,000 randomly-sampled pairs) correlates with cosine distance of the embeddings, Spearman r = 0.66 **(e)** Scaling the analysis up to all glycosyl hydrolase families and sub-families shows that there is variability in the agreement between phylogenetic distance and ESM2 embedding similarity. **(f)** GH1 family members coloured by their k-means cluster, projected from their embeddings into 2D using t-SNE as in panel C. **(g)** Adjusted Rand Index (ARI) and Normalized Mutual Information (NMI) between random seeds of k-means across various k-means, with error bars indicating one SD (based on 10 replicates). Three controls are shown: 80% subsampling (orange), embedding shuffling (red) and a distribution-matched control (based on a synthetic dataset with a matched data distribution, green) **(h)** Average sequence similarity within and between clusters in 2F, estimated using 1000 randomly sampled sequence pairs **(i)** (i) Distribution of protein aromaticity (percentage of residues that are aromatic) within and between the clusters from IF. (ii) Percentage of sampled families where the effect size of each feature was the greatest. Points represent individual features.

A very different challenge is to represent deep rather than widely spread sequences. Enzyme discovery does not only need to resolve sequences with different folds but should reveal a fine-grained structure within a family of enzymes with a single fold yet *different* activities.This is intrinsically a different challenge: single enzyme families are similar in sequence and structure but may harbour related activities in which their target substrates differ by as little as a single atom. One example of such a family is GH1, known to harbour both glucosidase and myrosinase activity, the key difference being the departure of an oxygen vs a sulfur. To investigate if this functional distinction is preserved in embedding space, we encoded members of this family of αβ-glycosyl hydrolases using MPEs. A functional distinction was again evident, with a clear cluster of myrosinases surrounded by β-glucosidases (Figure 2C). Other catalytic functions promoted by members of the family were found distinct in embedding space, with phosphoglycosidases occupying their separate regions.The observation that t-SNE can capture such functional distinctions suggests that embedding similarity in mean-pooled embedding space is a reflection of similar enzyme functions.

### Sequence similarity and phylogenetic approaches share mutual information with ESM2 MPEs within enzyme families but capture different evolutionary signals

We now aimed to benchmark how MPE similarities correspond to a known strategy for assessing protein relatedness – phylogenetic distance calculation based on sequence similarity. To assess the similarity of MPE embeddings to each other we used cosine similarity – another standard method for comparing the relatedness of a pair of embeddings^27^. Using cosine distance yields comparable results to Euclidian distance (Figure SI2iii, Spearman r = 0.99). When comparing MPE embedding similarities and phylogenetic distances between pairs of proteins, we would expect some correlation because evolutionary closeness is (albeit imperfectly) related to common function. For a random subset of GH1 enzymes, the relationship between the phylogenetic distance and the cosine similarity between embeddings is linear, but with many outliers where MPE similarity and phylogenetic relationship are not related (Figure 2D, Spearman r = 0.66). Changing to Euclidian distance impacts the correlation with phylogeny only marginally, even with L2 regularization of the embedding (Spearman r = 0.70/0.69 Figure SI2i and ii). Going on to test the generalizability of this finding, we studied every subfamily of glycosyl hydrolases in the CAZy database.There is variation in the strength of the relationship (as assessed by Spearman r) between these scores within diverse families of glycosyl hydrolases; for families 7, 15, 16_1 and 31 no correlation was discernible, while 2_8, 17, 137 and 193 exhibited near-perfect correlations with Spearman r > 0.94 (Figure 2E; Table S1).

The deviation from phylogeny in many cases supports the hypothesis that MPEs are capturing a different signal than evolutionary relatedness, or sequence similarity of enzymes^26^. Protein similarity calculated by MPEs used here correlates well with the sliced-Wasserstein embeddings (SWEs) used by Shaw et al.^16^ shown to compare functional similarity of proteins (Figure SI3i, Spearman r = 0.98), and MPEs correlated (albeit loosely) with sequence identity (Figure SI3iii, Spearman r = 0.77). Using the final eight layers of the embedding rather than the final layer only did reduce the correlations between SWE and MPEs (Figure SI3ii, Spearman r = 0.83), and the correlation dropped when substituting SWEs on the last 8 layers with MPEs for correlations with sequence identity (Figure SI3iv, Spearman r = 0.59). Given this good correlation between SWEs and MPEs, PLM-clust uses MPEs to represent proteins in a more computationally-efficient manner^16^.

### k-means applied to MPEs reproducibly captures sequence-function relationships

K-means is an unsupervised learning algorithm for data labelling that has been previously applied to protein language model embeddings for remote homology detection^28,29^. The k-means algorithm is based on the hypothetical case with roughly spherical distributions around a fixed number (i.e. ‘k’) centroids.The algorithm clusters data by adjusting the position of the centroids to minimize within-cluster sum of Euclidian squares. It can run with any number of specified centroids, and in its ‘hard’ formulation (used here), each datapoint is unambiguously assigned a cluster corresponding to its nearest centroid. Here we probe whether application of k-means on protein language model embeddings can suggest groups of functionally related proteins with diverse evolutionary origins, aiming to reach beyond the limits of detection of pure homology. To this end we first probe the robustness of prediction. To be useful, this approach should, as a minimum, make reproducible suggestions for clusters, independent of the random seed chosen. This property was thus tested by k-means labelling of GH1 MPEs resulting in the labelled t-SNE shown in Figure 2F. The clusters identified in high-dimensional embedding space form continuous regions when projected using t-SNE and are therefore consistent with the hypothesis that the clusters are capturing embedding-space neighbours.To test stability across random seeds for different clustering granularities, we applied k-means multiple times on the GH1 embeddings, varying the number of means to investigate how reproducibility is influenced by number of clusters. We found that both the Adjusted Rand Index (ARI) and the Normalized Mutual Information (NMI) are > 0.9 across replicates for k < 7 and drop down to a stable 0.65 as k increases to 20 (Figure 2G).This indicates that we have near-perfect run-to-run reproducibility for low values of k, yet there is a cluster threshold above which k-means can ‘overfit’ on embedding space, if too many degrees of freedom are provided, leading to stochastic outcomes. This approach provides an approach for selecting a number of means when clustering embedding space with k-means.

Given that the distance between embeddings must correlate to some extent with phylogenetic distance (which in turn relates to sequence similarity), we would expect a good clustering method to result in clusters that generally have greater internal sequence similarity than between each other.To test these assumptions, we calculated the intra- and inter-cluster mean sequence similarities using the k-means clusters from Figure 2F. The results are shown in Figure 2H. While some clusters have strong similarities within but not between, such as 1, 2, 3 and 5, some clusters show comparable similarity within and between, namely 4 and 6. Clusters 4 and 6 occupy central regions of the t-SNE and we interpret them as ‘bridging clusters’ across sequence and embedding space that also connect functionally related proteins (e.g. clusters 4 and 6 contain β-glucosidases and phosphoglucosidases catalysing the same chemical transformation on different substrates).

Now that we have a reproducible and functionally meaningful method to identify clusters in embedding space, we can examine similarities and differences in the sequences within and between clusters to identify the biophysical drivers of similarity in embedding space.To do this, we add features to each sequence in the dataset with a panel of biophysical labels and then run statistical tests to identify features whose values vary systematically between clusters.We quantify the ability of a biophysical trait to resolve clusters using ANOVA effect size. For the model case of GH1 family of proteins, when separated into 6 clusters, cluster identity is driven by differences in proline content, amino acid polarity and predicted structural flexibility. In contrast, other features, such as the presence/absence of signal peptides and helix fraction, are not features that are well-resolved between the clusters (**Figure 2i(i)**).This analysis suggests that biophysical features determine closeness in embedding space. Scaling up experimentation to cover other GH families, we find that feature importance varies depending on the specific family – no single feature is consistently the primary determinant of clustering (**Figure 2i(ii)**), indicating that different molecular recognition motifs drive embedding distribution between families.This observation is consistent with the hypothesis that using a foundation model (even without fine-tuning) can effectively separate members of a single enzyme family to refine CAZY’s organisational category families with higher resolution, as the model has implicitly ‘learned’ features that drive functional differences within families.

### Pseudo-perplexity estimation from raw logits enables rapid, generalizable fitness scoring

Efforts to reduce the computational burden of fitness scoring with protein language models, which scales with the length of the input, have focused on finetuning the outputs of masked language models with a helper model^30^ (also applied to ESM2^31^) or evaluating them. To this end *pseudo-perplexity* (PPL*) is normally calculated for an input by masking tokens one-by-one and then calculating the likelihood that the model’s prediction matches the ground-truth, averaged over all sampled tokens, and has been shown to be effective for estimating masked language model performance^30^ (Figure 3Ai). However, to make PPL* calculations runnable at metagenomic scale, a speedup of several orders of magnitude is required to make the computation feasible. A second evaluation strategy involves masking of a certain proportion of the sequence, typically 15%, instead of one-at-a-time masking in masked language models in general^32^ and also specifically for protein sequences^13,33^ (Figure 3Aii). We take the premise established in previous work^31^ that information on the masked likelihood of a token is encoded in the forward pass of that unmasked token one step further, and test the approximation that it is in fact linearly encoded and does not need further processing other than linear extraction of the logits for the true token, requiring no bespoke treatment of embeddings and therefore accelerating PPL* calculations.

**Figure 3:**
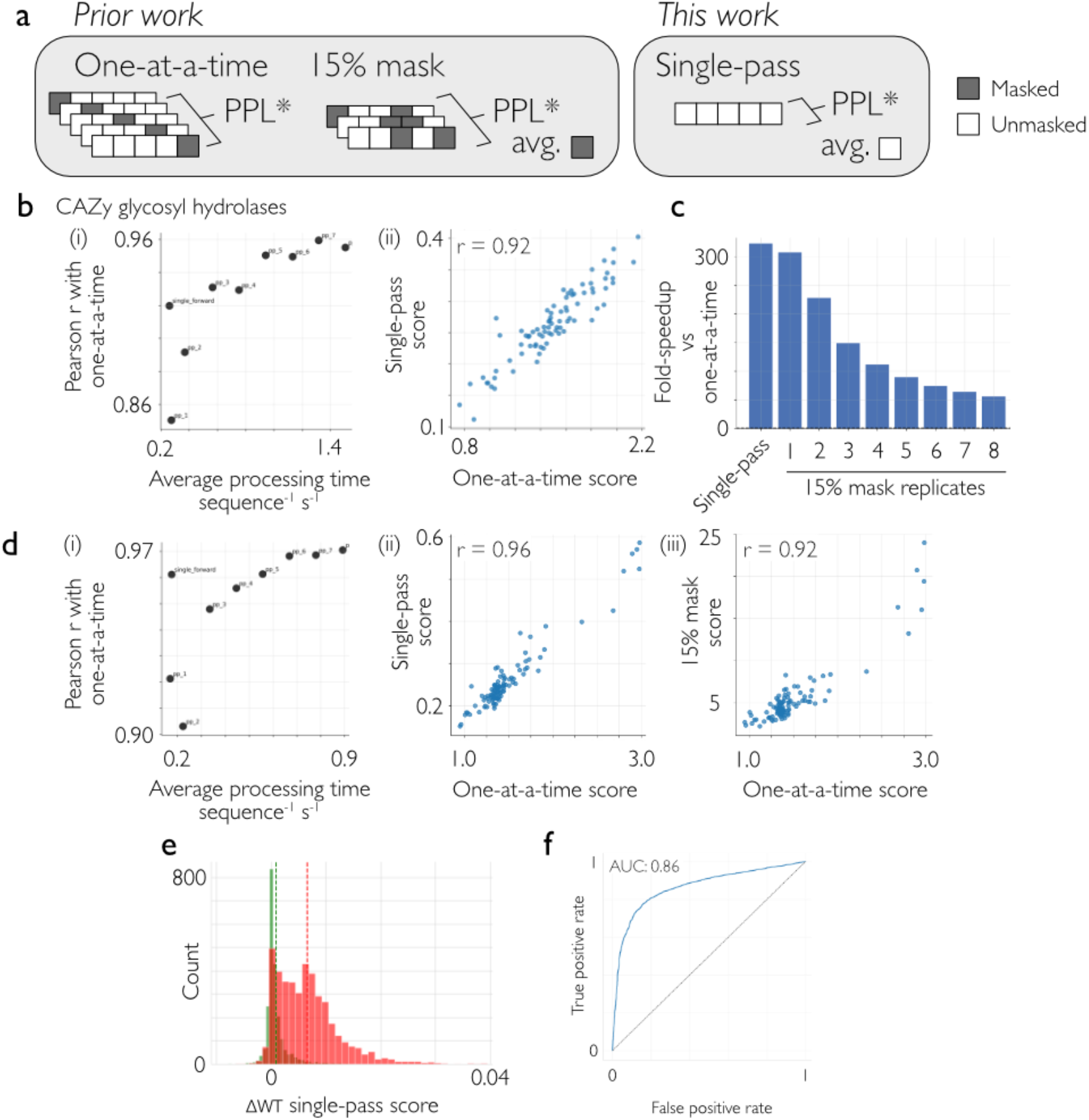
Accelerating accurate PPL* estimation using a single-pass method. **(a)** Methods available to generate PPL* scores from masked language models like ESM2 to reduce the computational burden of PLM-clust: (i) one-at-a-time masking, where each residue is masked in turn and the average cross-entropy between the correct amino acid and the model’s estimation is calculated (ii) 15% masking, where 15% of the residues are masked per pass, and the average cross-entropy between the correct amino acids and the model’s estimation is calculated. Multiple replicates of the random masking process can be carried out and their results averaged to account for the randomness introduced by the masking. (iii) single-pass (this work), where no masking is carried out, and the average cross-entropy between the true sequence and the model’s estimation is calculated. **(b)** One member of each glycosyl hydrolase family was selected at random, resulting in >100 datapoints, and each method for PPL* estimation was carried out.The 15% masking method was carried out with replicates between 1 and 8. The correlation between each method (the 8 15% methods and the single-pass method) is plotted in (i) against the one-at-a-time score for each sequence.The underlying data for the single-pass relationship with one-at-a-time is shown in (ii), where each point represents a member from a separate CAZy family. **(c)** The relative speed-up of each method compared to the one-at-a-time, measured in fold-decrease in processing time across the dataset in (b). **(d)** The same analysis as in (Bi), but for 100 randomly-selected homologues of the same glycosyl hydrolase 11 (GH11) family, which share the same fold. (ii) the correlation of single-pass and (iii) of a single 15% mask with one-at-a-time for the dataset in (Di). **(e)** Random sample of 6421 protein-mutant pairs from the proteingym clinical substitutions dataset scored by change in PPL* from their ‘wild-type’ sequence – higher score indicates model predicts lower fitness. (F) ROC curve for the dataset in E – AUC of 0.86, classification of the full dataset into ‘pathogenic’ or ‘benign’.

As one-at-a-time masking is the gold-standard^34^, we set out to benchmark the correlations between the 15% masking approach or single-pass PPL* and that deriving from one-at-a-time masking. We tested a range of diverse sequences and structures deriving from the glycosyl hydrolases of the CAZy database, selecting a member from each GH family at random. To account for the randomness inherent to the 15% masking approach, we performed up to (and including) 8 random 15% masks/sequence, as the random nature of the masks can be accounted for with multiple maskings followed by averaging.We find that all speed-up methods correlate well (Spearman r>0.87) with one-at-a-time masking (Figure 3Bi). The worst trialled method was a single 15% mask (Spearman r = 0.87), but this score increased to a plateau of Spearman r = 0.96 after 7 replicates of the random masking process. In comparison, the single-pass method achieved a Spearman r = 0.92 compared to the one-at-a-time masking process (Figure 3Bii). On the sequences used (limited to length <500 aa), the speedup of single-pass was 320x compared to the one-at-a-time (Figure 3C). The speed of the single-forward pass was between the single and double 15% masking replicates, with a similar Spearman r (vs one-at-a-time) to the double 15% masking (Spearman r = 0.91) (Figure 3C).

We have shown that this accuracy is retained between distantly related folds of protein, but we must also ask if the same correlations hold for more closely-related sequences, such as those within a single fold – in other words, are the methods precise enough to resolve differences between proteins with high sequence identity? To answer this question, we re-ran the analysis on the GH11 family of endo-xylanases, which all inhabit the same jelly-roll fold structure and have similar catalytic functions to each other^35^.We drew comparable conclusions to the diverse space search performed before (Figure 3D).

We conclude that all trialled methods are suited to substitute the slow one-at-a-time masking when calculating the PPL*s of diverse protein sequences using the ESM2 model. Single-pass processing has the further advantage that it is deterministic (it will not change between runs as there is no random mask), and can be calculated in the same forward pass that is used to generate the MPEs that are used for embedding clustering, affording it a further 2-fold speedup compared to passing the sequence through the model once for clustering and once for scoring. Single-pass processing affords a multi-order-of-magnitude speedup of the PPL* calculation, decreasing processing time from >50h on consumer-grade equipment to less than half an hour for a typical dataset. Now, we may ask whether single-pass PPL* thus generated report well on protein benchmarks of fitness. Using proteingym^36^ clinical substitutions dataset, we find that pathogenic mutations have significantly worse scores derived from single-pass PPL* than do neutral ones (Figure 3E), with a classification ROC-AUC of 0.86 and an average precision of 92% (Figure 3F).This suggests that the single-pass PPL* metric is a meaningful representation of the ‘functionality’ of a protein.

### Case Study 1: Identifying new functions within a well-annotated enzyme family

Xylan is a linear polymer with sidechains, abundantly present in plants and important in the food industry to improve the taste and material properties of baked and cooked goods. However, to achieve optimal effects, xylan should sometimes be processed into shorter, more soluble xylo-oligosaccharides (XOSs) and a small amount of free xylose. The short XOS enhance food matrix interactions^37-39^, while the xylose promotes browning of the food by the Maillard reaction^40^.To produce XOS, the GH11 family of xylanases is of keen biotechnological interest,^41^ cleaving internally in the xylan polymer. However, GH11 is known to harbour a strict molecular recognition requirement for longer xylan chains for hydrolysis to occur, resulting in low yields of the monosaccharide from polymeric xylan^42-44^. This means that other helper enzymes or monosaccharide additives must be added to promote browning. We proposed a GH11-only strategy, which involves identifying GH11 enzymes that break the paradigm of the ‘true xylanase’ and release usable amounts of free xylose alongside the XOS products, simplifying the required enzyme cocktail and increasing the scalability of the process on the biocatalytic level^45^. This means we are looking for promiscuous xylanase/xylosidases as bi-functional enzymes (Figure 4B). To uncover hidden promiscuity in the existing GH11 corpus, we applied PLM-clust to the problem. As we were searching for thermostable enzymes, we anticipated that using PPL* was also likely to reveal good candidates for the biocatalytic process.

**Figure 4:**
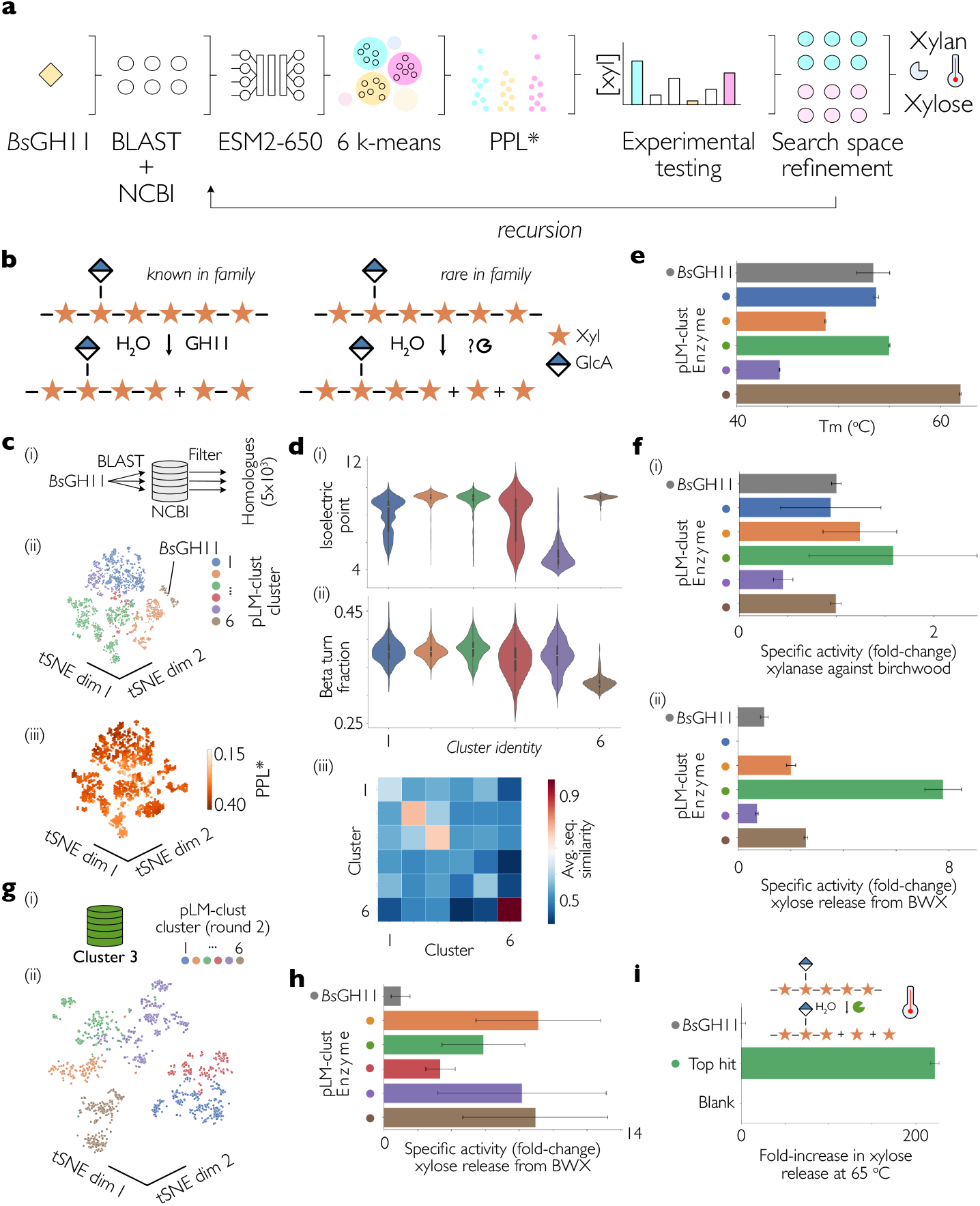
Discovery of untapped catalytic potential in a well-known enzyme family (case study 1) (a) The pipeline for iterative clustering and annotation of the embedding space of ESM2. (i) A natural sequence corpus is gathered based on a retrieval tool such as BLAST, based on an ‘anchor’ in sequence space (ii) The sequences are passed forwards through ESM2, and the final embedding layer is extracted, and subsequently collapsed, making all embeddings the same 1024×1 dimension, independent of protein length. (iii) Embedding space is segmented into N segments using a hard k-means algorithm, where N is the experimental capacity of the round. (iv) Sequences are ranked according to the average NLL of their unmasked identity and top hits are promoted to the experimental pipeline (v) Multi-trait experimental measurements are carried out, reporting on stability, activity and promiscuity of the promoted sequences (vi) The embedding space with the highest experimental activity is re-entered into the pipe at position (ii). (b) Known activity in the GH11 family of true xylanases, alongside the target activity that we set out to look for. (c) Example implementation for the GH11 family of xylanases. (i) BsGH11 is used as a query sequence to the NCBI nr database to retrieve 5000 sequences. (ii) The embedding space of GH11 is not continuous but split into regions that are connected to a varied degree. k-means with 6 means is applied in the high-dimensional space, whose structure is sufficiently preserved when projected onto two dimensions so that the identified clusters still appear meaningful. (iii) When scored, ESM2 does not evenly score sequence space, but displays preference for some regions more than others. This means that segmenting the space and then picking the top score from each cluster is critical to maintain diversity. (d) Selected biophysical parameters distributed between clusters as defined in (Cii), and average sequence identity within and between clusters for a random sample of 1000 protein pairs. Apparent melting temperature as measured by thermal shift assay (TSA). Error bars represent the standard deviation of three independent replicates. (f) Relative specific activity of cluster representatives against the substrates (i) azo-birchwood xylan and (ii) beechwood xylan relative to BsGH11. For (ii), maximal rate of xylose release is quantified. Error bars represent the standard deviation of three independent replicates. (g) Recursive mining of sequence space, using cluster 3 as the search space for the next round of discovery. (ii) t-SNE of embedded cluster 3 with 6 k-means clusters assigned. (h) Relative specific activity of xylose release from beechwood xylan of each of the cluster representatives, compared to BsGH11. (I) Relative release of xylose from a slurry of beechwood xylan over a 2h incubation at 65 °C.

Using a canonical GH11 enzyme *Bs*GH11 as bait^46-49^, we retrieved, embedded and clustered 5000 sequences from NCBI into 6 clusters.The embedding space was found to produce some discrete clusters, and other regions of more well-connected embedding space (Figure 4Cii). This indicates that some proteins have very similar embeddings that are very different from all other embeddings, while other proteins have embeddings that form smooth bridges in the space. *Bs*GH11 joined a discrete cluster (as per t-SNE projection, commonly used in visualizing language model embeddings^18^). Scoring these sequences using PPL* resulted in the labelled t-SNE shown in Figure 4Ciii, and shows that scores are not randomly distributed through the space, with some regions having systematically higher or lower scores. This highlights the pitfalls of selecting enzymes based on zero-shot scoring without clustering to enforce diversity – model phylogenetic biases^50^ mean that only a very constrained sequence space would be sampled (the top 34 PPL* scores belonging to members from clusters 1 and 5).

Using the biophysical interpretability framework established in Figure 2 to open the ‘black box’, we investigated the GH11 clusters to identify the features driving clustering. We found that some clusters carried distinct biophysical profiles that are identified by the MPEs but not sequence similarity – for example, cluster 5 members carried an isoelectric point of pH 5, compared to other clusters which had a pI of on average 8 (Figure 4Di). Despite having distinct pI profiles, clusters 1 and 5 are not resolved by sequence similarity (figure 4Diii) – but k-means in high-dimensional embedding space was able to resolve this stark biophysical difference. Despite some clusters bearing obvious features that mark their identity, most clusters could not be assigned one outstanding biophysical feature, although some appeared to be correlate high sequence similarity (Figure 4Diii).This suggests that clustering is in some cases driven by a composite of features not detected by our biophysical panel, or sequence similarity alone. A full account of features and their effect sizes is found in Table S2.

To examine how these distinct profiles translate into catalytic performance, we synthesised the top-scoring candidate from each cluster.We achieved high (>10 mg/L) purified yield of 5/6 of the cluster representatives – the non-expressing member (cluster 4) belonged to a cluster enriched in truncated protein sequences, making it likely that this cluster contained only non-functional sequences (Figure S5). This failure serves as a guide for using PLM-clust – manual inspection of sequences before proceeding would have detected the obviously truncated sequence. All expressed enzymes were found to be proficient endo-xylanases on birchwood xylan (Figure 4F), with between 0.4-1.6-fold the specific activity of the reference BsGH11 enzyme, indicating that all clusters were able to recognise and cleave the xylan polysaccharide. For reference, empirically 49% of recombinant proteins are expected to be expressible and 52% of these purifiable^51^.

Measuring the xylose release from beechwood xylan, we identified that the enzyme from cluster 3 had an 8-fold greater maximum rate of release compared to the wild-type *Bs*GH11 (Figure 4fii). In contrast, although the enzyme from cluster 1 had 0.95-fold the specific activity of the *Bs*GH11 reference enzyme at the endo-cleavage challenge (Figure 4fi), it had no detectable xylose release under the assay conditions.

### Recursion to navigate embedding space with unsupervised learning

Now that our embedding space contained functional labels, we targeted the search to the productive cluster 3 to investigate where maximal xylose release could be found in embedding space. We applied the methodology in Figure 4A recursively, re-clustering and scoring the members of cluster 3, and synthesising the top cluster members, with the results of the re-clustering shown in Figure 4G.

Four out of the five members were cloned, expressed and purified well; cluster member 1 was lost during the cloning stage, and was not pursued further. Further improvements in melting temperature were seen, with the best melting temperature (H9, iteration 2, cluster 3, green) being 69 °C, an increase of 17 °C compared to the enzyme starting point (Figure S6). H9 also exhibited a 6-fold increase in maximal xylose release rate / mg enzyme compared to *Bs*GH11. (Figure 4H) Indeed, all assayed second-round variants were able to release xylose from BWX faster than the reference *Bs*GH11, demonstrating that the embedding space of cluster 3 was indeed enriched in the function of interest.

A good enzyme candidate for process scale-up would have, amongst other properties, a high melting temperature, specific activity and high yield of purified protein. H9 (green) had a good combination of these properties according to our *in vitro* screening, and so we moved to a small-scale reaction under slurry conditions, with beechwood xylan present above its solubility limit in mildly basic solution (30 g/L, pH 8.0). We compared the xylose content of the supernatant of the slurry after two hours treatment with H9 compared to *Bs*GH11 at 65°C and indeed detected appreciable xylose release with H9 but not with *Bs*GH11 – over 200-fold increased release of xylose (Figure 4i), representing final xylose content of 0.3% w/w after two hours of reaction with the discovered enzyme H9, compared to non-detectable levels in *Bs*GH11. *Bs*GH11 was poorly suited to the task due to its low activity and relative thermal instability. Compared to mutational stabilization of *Bs*GH11, which was only able to increase the Tm by 2 °C^48^, our 17 °C increase in T_m_ and simultaneous improvement of catalytic features supports embedding space navigation with zero-shot scoring as an alternative to mutagenesis of a single starting point for finding enzymes for a target process. The evolutionary time spanned by the PLM-clust algorithm was on the order of hundreds of millions of years. A calibrated phylogenetic tree of the *Bacillales* members (representing only 15% of the total sequences studied here, which included representatives from both fungi and bacteria) suggested that the cluster representative in our first-round query *Bs*GH11 cluster was 101 million years (CI: 41, 157) away from *Bs*GH11 (SI Figure 5).

**Figure 5:**
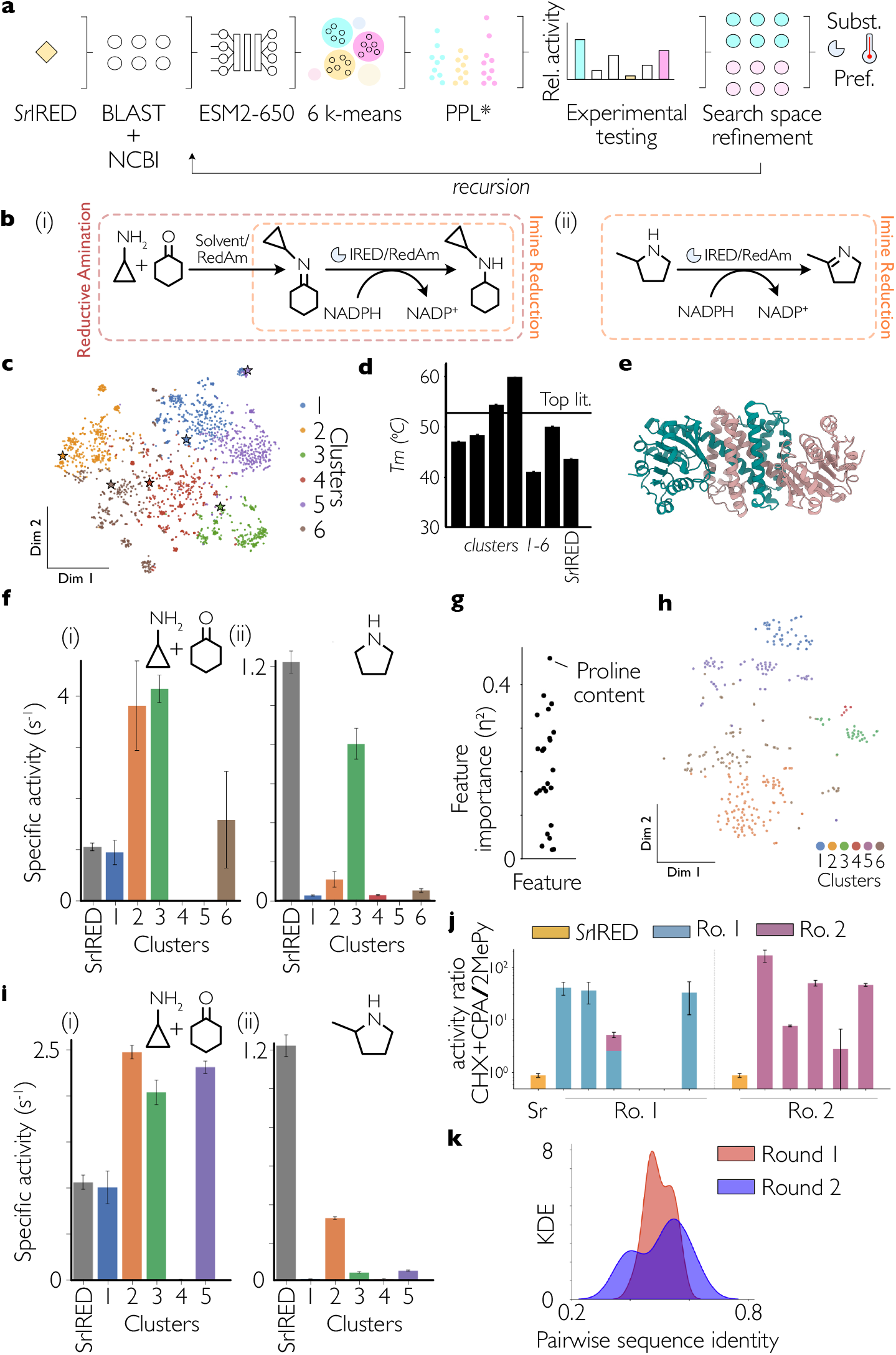
High-success enzyme discovery of promiscuous, thermostable enzymes for chiral amine synthesis. **(A)** Pipeline used to build the IRED input dataset and run PLM-clust. **(B)** Model substrates (i) Cyclohexanone (CHX) + Cyclopropylamine (CPA) and (ii) 2-Methyl-1-Pyrroline (2MePy). **(C)** Clustered IRED embedding space, annotated by k-mean cluster identity, shown by t-SNE. **(D)** Thermostability of sampled IREDs, as measured by thermal shift assay. **(E)** Alphafold3 predicted structure of thermostable cluster representative 3, coloured by monomeric unit. **(F)** Specific activity profiles of enzymes against (i) CPA/CHX (ii) 2MePy. **(G)** Feature importance of each of the sampled biophysical features for the IRED clustering. **(H)** Re-clustering and labelling of the previous cluster 3. Stars indicate the experimentally tested members of each cluster **(I)** Specific activity profiles of second-round enzymes against (i) CPA/CHX (ii) 2MePy. Error bars represent 1 SD of three independent reactions. **(J)** Relative specific activities (log-scale) of mined enzymes towards CPA/CHX divided by 2MePy. Error bars represent 1 standard deviation. Ro. 1/2 = round 1, Sr= SrIRED. Cluster member 3 from recursion 1 is bi-coloured as it is the founding member of the second round. **(K)** Kernel Density Estimation (KDE) of sequence pairwise identities within the first round and second round of screening.

### Case Study 2: Identifying promiscuous enzymes as starting points for directed evolution

Synthesis of chiral amines is a pressing pharmaceutical challenge, due to their importance in the production of many high-value chemicals^52^.These bonds can made from pre-formed imines by the imine reductase (IRED) enzyme class, of which hundreds of members have been characterized.^53^ Closely related is the reductive aminase (RedAm) enzyme class, which show a preference for binding to amine/ketone pairs and forming the imine intermediate *in situ*.^54^ They have advantages over IREDs, as they do not require pre-formation of the imine in solution, which is otherwise driven by the thermodynamic equilibrium in the solvent used (Figure 5B). However, distinguishing IREDs from RedAms is not trivial from sequence alone.^54^ Using substrate promiscuity for an amine/ketone substrate pair Cyclohexanone/Cyclopropylamine (CHX/CPA) as compared to a preformed imine 2-Methyl-1-pyrroline (2MePy, Figure 5B), we set out to identify if PLM-clust could be used to discover novel thermostable RedAms from the sequence corpus.Analogously to the hydrolase case, we were also targeting high thermal stability. As a reference enzyme, we used the fungal IRED *Sr*IRED, which is known to have promiscuous activity against CHX/CPA and 2MePy^55^. Applying the pipeline described in Figure 5A, we clustered the diversity of IREDs/RedAms (Figure 5C) and selected the top 6 cluster representatives for experimental characterization. All six of the enzymes expressed in high yield (>10 mg/L of culture) and had melting temperatures between 41-60 °C (Figure 5D). This included two IREDs that showed the highest reported thermal stability of any previously reported wild-type enzymes (only 4 °C below the top published thermostability-*engineered* IRED^56^).As before, PLM-clust therefore delivered a maximal hit rate (100%) of well-expressing, stable proteins, outstripping SSNs for sequence space exploration^5^. The diversity of enzymes screened was high, with a maximum sequence identity of 62% and a minimum of 43% (SI Figure 5), suggesting that novel sequence space was explored. The predicted structure of the most thermostable cluster member (from cluster 4) is shown in Figure 5E.This suggests assigning function is possible in low homology context, i.e. that PLM-clust can penetrate sequence contexts that homology searches would not have been confident to assign functionally.

We now set out to classify the catalytic features of each cluster. Four out of six clusters were active against CHX/CPA (Figure 5Fi), while five out of six were active on MePy (Figure 5Fii), leaving only cluster 5 with no identified IRED activity. Cluster 3 had high promiscuous activities against the model substrate pairs tested (Figure 5Fi and ii), and a high melting temperature (54 °C). Proline content was the most defining differentiator between clusters, as quantified by effect size, but many other features such as amino acid composition and pI also had meaningful effect sizes, making it hard to identify a single biophysical feature that best described the clustering and suggesting a more non-linear reasoning for clustering (Figure 5G).A full account of features and their effect sizes is found in Table S3. Cluster 3 was used as the search space for the second pass of PLM-clust, with the goal of identifying additional stable enzymes with RedAm-like promiscuity profiles.

Following embedding, scoring, and synthesis of the cluster representatives for this sub-search-space (Figure 5H), all 5 of the new cluster representatives expressed with good yield (>10 mg/L culture) with melting temperatures between 44 °C and 55 °C (SI Figure 6). Four of the five new enzymes displayed activity against both model substrates (Figure 5I). The founding member steers the predictions: their substrate specificity (measured as the ratio of rates for substrates CHX/CPA vs 2MePy) is increased in the mutants derived from it (Figure 5J). In a major deviation from purely sequence-based clustering, the average pairwise identity of enzymes in round 2 did not fall compared to that of round 1 (Figure 5K). This is despite the enrichment of enzymes with promiscuous activities against the pair of substrates used.

**Figure 6:**
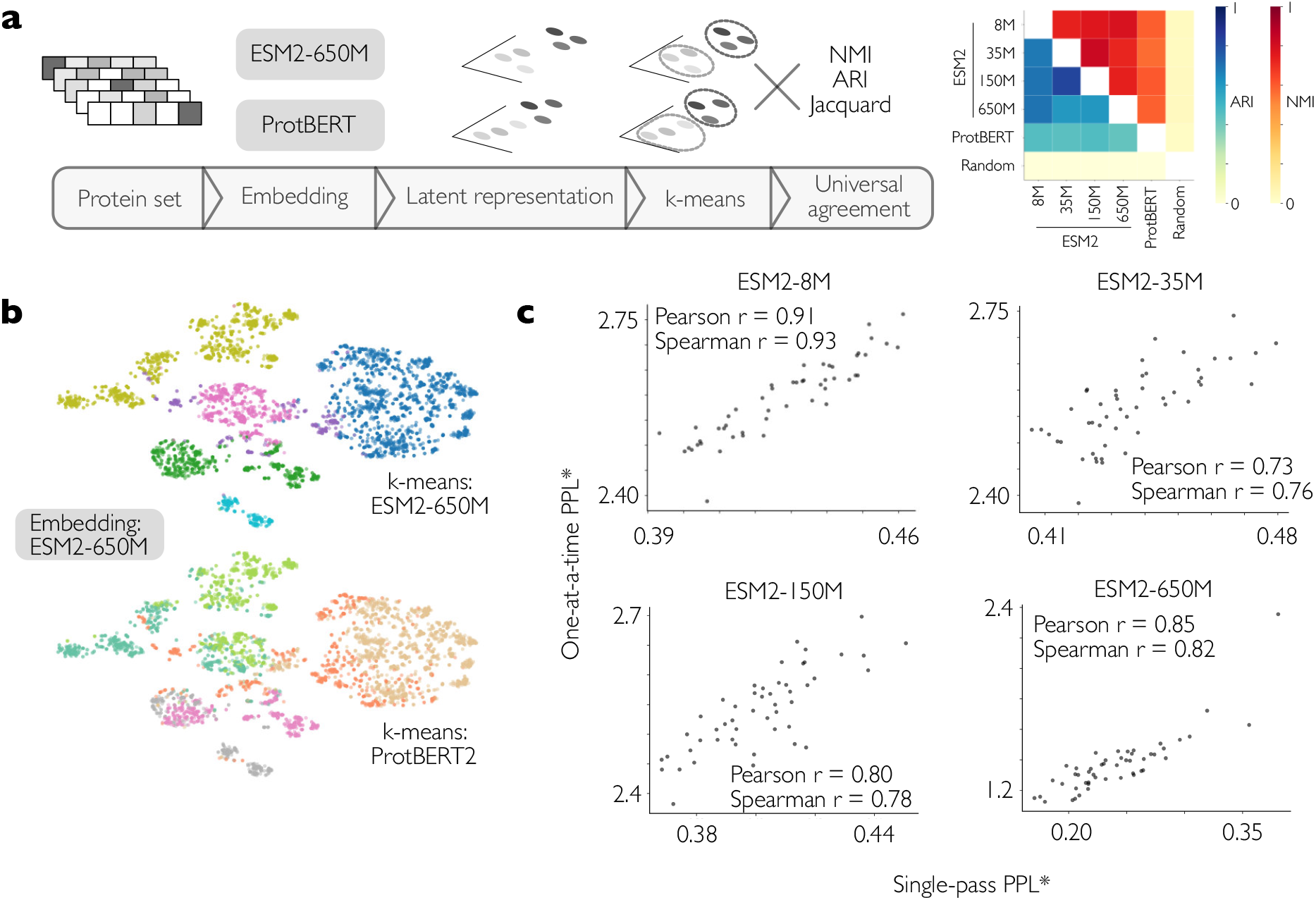
PLMs arrive at comparable representations of proteins in their embedding space, and these representations are consistently extracted by PLM-clust. **(A)** Left, framework for comparing PLM-clust results between language models. Right, Adjusted Rand Index (ARI) and Normalized Mutual Information (NMI) between the clusters derived from different PLMs following PLM-clust. The negative control used a feature-shuffled embedding null, where each of the embedding dimensions of the ESM2-650M-embedded sequences were shuffled between sequences, before re-applying k-means. Sequences are the GH11 iteration 1 set from Case Study 1. **(B)** (Top) Sequences used in (A) embedded using ESM2-650M, MPEs calculated and visualized using t-SNE. Colouring is based on cluster identity, assigned using PLM-clust on the ESM2-650M embeddings, with 6 means. (Bottom) as above, but colouring is based on PLM-clust on the ProtBERT embeddings, with 6 means. (C) Comparison of the PPL* scores calculated using single-pass and one-at-a-time strategies for a random set of 50 of the enzymes in (B) using four different PLMs.

### PLM-clust is a generally applicable framework for linking protein foundation models to enzyme discovery

The principle of embedding clustering using unsupervised learning is theoretically applicable to any used protein language model. However, it is not clear if the predictions from different models would converge or diverge. This is because each model has different training data, different internal architecture and a history of training with different objectives. To test the convergence of predictions from existing protein language models, we applied a set of different masked language models to a set of enzyme orthologues (those used in our xylanase discovery campaign, Figure 6A). Using 6 k-means centroids, we established that there is significant mutual information between k-means clusters on MPEs from four ESM2 models (8M, 35M, 150M and 650M) and ProtBERT^57^ (Figure 6A). These masked models are comparable as they are all trained with the same functional objective – masked sequence prediction given some context, although they have architectural differences^13,57^. To visually depict the agreement in clustering and cluster assignment between models, we visually overlayed the cluster identities from PLM-clust run on ESM2-650M with representations of the same proteins generated by ProtBERT (Figure 6B). We see that the clusters calculated using ESM2-650M are coherent with the representations generated by ProtBERT. The finding that PLM-clust comes to comparable conclusions when bolted to multiple language models supports the following hypothesises: 1. MPEs are information-rich strategies for comparing protein language model embeddings regardless of chosen model 2. Modern protein language models represent proteins in ways that converge in their representations of protein similarity 3. K-means performs consistently well on unregularized final embedding layers for identifying these similarities.

The second feature of the PLM-clust pipeline is single-pass PPL* estimation. To identify if this strategy is valid for models other than ESM2-650M, we ran correlations of the single-site method against the single-pass method for four other PLMs. We found correlations between the results between Spearman and Pearson r of 0.7 and 0.9 for the used dataset of functionally similar proteins (GH11s) and diverse proteins from the CAZy database (SI figure 7, Pearson, Spearman > 0.88, 0.87 for the ESM families of models, Pearson, Spearman = 0.70, 0.70 for ProtBERT).The excellent correlation coefficients suggest that single-pass PPL* estimation can accelerate zero-shot scoring of diverse proteins for all the PLM used. The finding that both clustering and scoring using PLM-clust is valid between multiple masked protein language models underlines that the functional assignment of unknown proteins is successful regardless of the specific model. As new models become available, PLM-clust will be an evergreen framework that can be applied generally to enzyme discovery campaigns, mirroring and improving upon the use of SSN-linked protein homology in protein discovery campaigns.

## Discussion

Using the embedding space of a protein language model breaks the reliance on protein sequence homology for predicting functional relatedness between proteins. Here we exploit this to direct protein foundation models towards identifying sequence-diverse enzymes to annotate them reliably with function and improve their properties compared to the starting points. We achieve this by using recursive clustering and only require experimental confirmation on a small number of suggested datapoints. In contrast to directed evolution of a known starting point (diversified e.g. by error-prone PCR or site saturation mutagenesis) and following the prevalent ‘one amino acid at a time’ paradigm^58,59^, we are able to rapidly traverse sequence space down to less than 50% homology between candidates per round. Detecting isofunctional enzymes over such sequence distances goes beyond homology predictions, may lead to different mechanistic features and evolvability, and is also sufficient to overcome patent protection (typically set at 60% homology to a protected sequence). Experimentally our testing of candidates achieves orders-of-magnitude greater hit rates than typical for directed evolution workflows,, while covering a wider perimeter of sequence diversity (<50% homology). Such diversity corresponds to hundreds of millions of years in notional evolution time to reach as far as <50% homology^60^) and even withing clusters diversity corresponding to 100 million years is observed (SI Figure 5). We obtain much improved properties (up to 6-fold specific activity, 17 °C in thermostability in one case-study that led to two orders of magnitude improvements in target activity under conditions too extreme for the reference enzyme) with only 10 candidates tested per campaign in two batches, a small fraction of the effort of an experimental evolution campaign that typically involves screening of >10^3^ mutants or more. Compared to EVOLVEpro^61^, the state-of-the-art active learning approach for protein engineering incorporating pretrained language models, we required only two rounds to achieve improved mutants (compared to between 4 and 7).This is likely because we propose existent enzymes instead of the riskier variant-proposal approach taken by EVOLVEpro, but may also depend on the target enzyme class.The same applies to the MULTI-evolve strategy^62^, which aims to extract epistatic interactions using pretrained models and data collection on mutants around a wild-type. MULTI-evolve was successful in reducing the number of rounds to 3, but is still inherently constrained by the local epistatic landscape of the chosen starting point, and at the expense of testing over 100 target mutants to obtain sufficient training data.

We show that making effective use of natural sequence space can be at least as targeted to the task of identifying improved enzymes as state-of-the-art directed evolution^61^.This may be because protein foundation models have learned useful representations of neutrally (i.e. non-adaptively) drifting proteins, that can be exploited to relieve local epistatic constraints on functional adaptation^6^. This neutral drift has long been postulated to support neofunctionalization^63^, and indeed much of the signal in metagenomic datasets may report on this neutral drift, rather than positive Darwinian selection^64^.

Our strategy requires no model fine-tuning,^20^ which makes it possible to use open-source foundation models directly off-the-shelf. This is a distinct advantage as fine-tuning or explicit training^65^ of protein language models to predict function has so far met with at best mixed or downright unreliable results: in several well-documented examples fine-tuning has failed to achieve meaningful performance gains,^66^ or has even decreased performance. In contrast, our evidence for consistently high hit-rates (second round > 90%) predicting well-expressing, functional enzymes and orders-of-magnitude improvements substantially reduces the risk associated with enzyme discovery. We use a simple mean-pooling of the final embedding layer, and show that it results in protein similarity calculations comparable to theoretically more information-rich methods such as SWE^16^.

We show that two rounds of computation-experimentation allows for implicit specification of target reaction conditions, which are then applied to the foundation model embeddings via unsupervised learning algorithms. We show that these applied conditions can be complex, starting with the ability to catalyse a target reaction but also including catalytic promiscuity^67^, operating temperature and pH. This ability is in contrast to the state of the art in functional fine-tuning of protein language models for enzymes, where ‘function’ is often reduced to a single E.C. class^65^ or semi-automated database tags^60,68^.

While language models like ESMFold have been used to predict 3D structure, function cannot straightforwardly be read from structure^69^. Predictions that extrapolate from structure must therefore include experimental validation. However, such validation has been missing e.g. in a language model-based prediction of E.C. numbers,^69^ questioning whether assignments are premature^70^. Our evidence suggests that PLM-clust is able to extract the information needed for enzyme discovery from pretrained foundation models using a low amount of test data (obtained under the conditions of interest) providing a powerful alternative to finetuning models on target functions. The success of PLM-clust is based on the availability of metagenomic data, both for the initial foundation model training, and for iterative searching. Therefore, we argue that the continued collection and sequencing of environmental DNA (eDNA) will continue to power enzyme and protein discovery in the future, even as directed evolution becomes automated and move *in silico*.

The agreement of MPEs derived from ESM2 with phylogeny demonstrated in this work introduces the possibility that enzyme discovery via ancestral sequence reconstruction (ASR) could benefit from integration with PLM-clust. Enzymes discovered by ASR tend to have greater thermostability than their extant counterparts^71^, with tuneable substrate preference^72^. ASR involves creating a larger number of candidate sequences than can be screened – applying PLM-clust could assist with candidate selection for experimental work. PLM-clust may be used to identify functional ‘swapping over’ points in the continuous phylogeny of enzymes and therefore streamline candidate enzyme testing.

Straightforward practical application - via the pipeline codebase or a python package (via github, see below), directly through our web portal (discovery.plmclust.co.uk) or via a web-hosted Google Colab notebook - means that researchers can build the framework into existing discovery pipelines, e.g. in larger scale enzyme annotation of billions of unknown ORFs contained in large databases such as MGnify^73^. Ultimately easier access to *bona fide* enzymes with predictable, improved function will make more enzymes available for use in sustainable biocatalytic processes or as biological precision reagents.

## Methods

### Online methods

#### Google Colab implementation

The code is available to run in a user-friendly way via a Google Colab workbook (SI Figure 8). Here, users can cluster sequences of interest with a configurable number of clusters (k), and visualize the results using dimensionality reduction.Top k sequences are proposed to the user for experimental validation. A csv of all sequences and their associated clusters is available, along with their calculated PPLs.The user can also implement a silhouette score analysis to estimate the optimal number of clusters in the data, without relying on pre-defined k-means. Access at: https://colab.research.google.com/drive/1FzRK-jEWgY4MOr6c9Os8MCXzZ4KSdEWA?usp=sharing

#### Python package

Users can install the PLM-clust code via pip, using pip install uht-discovery. This also installs accessory modules that can be used for automated BLAST search of NCBI’s servers, perform quality control of sequences, and run clustering. The package has a CLI and a GUI that users can interact with or embed in larger data processing pipelines. Code has been optimized to run on Apple Silicon metal performance shaders (MPS). The package repo can be found at https://github.com/Matt115A/uht-discovery-package.git

#### Web-server

Users may submit jobs to the PLM-clust web server (plmclust.co.uk) – a user may start with a single sequence, retrieve homologues *via* BLAST of NCBI’s databases, and then run quality control followed by clustering with a configurable number of k-means centroids.The resulting t-SNEs and UMAPs can be viewed interactively, and cluster representatives are downloadable from the web server.

#### Acquisition and curation of benchmark datasets

The CAZy database was downloaded from www.cazy.org, and the Uniprot accession codes were used to retrieve sequences from the UniProt database *via* the API. Proteins were identified as either glycosyl hydrolases (GH) or glycosyl transferases (GT) using exact string search for either ‘GH’ or ‘GT’ in the sequence header. Proteins filtered for length, removing sequences over 5000 amino acids. An exact string search for CAZy family annotations was used to separate the list of sequences into separate CAZy families, and each family was filtered to remove any sequences with > 1 standard deviation in length from the median length. This focused the datasets on sequences with consistent and comparable domain structures (the dominant one in each of the datasets).

#### tSNE representations of CAZy and GH1

Following running PLM-clust on the CAZy sample as derived above and t-SNE generation, sequences with only ‘GH’ or ‘GT’ in their sequence header were coloured and displayed.The input file is Extended Data 9. For GH1, the CAZy database was specifically subsampled, using sequence retrieval of all GH1 enzymes with known annotated function, and a random sample of 1000 non-characterized GH1 enzymes with NCBI accessions in the CAZy database were also downloaded. In total, this resulted in 382 characterized sequences, and after manual length filtering to remove multi-domain enzymes, a total of 1194 (characterize and uncharacterized) sequences remained. Sequences were then coloured according to their labelled E.C. class.

#### SSN construction

The EFI-EST webtool^74,75^ was used to construct an SSN given the input set of sequences used for the CAZy database benchmarking (Extended Data 9). As recommended, alignment cutoffs were selected based on inflection points in the number of edges when compared to the alignment score threshold. The parameters used to generate the network are available in Extended data 10. The network was visualized in Cytoscape using a Perfuse Force-Directed Open CL method.

#### Embedding protein sequences using MPEs

Sequences were embedded using ESM2 (ESM2_t33_650M_UR50D) on MPS, and the final embedding layer ([33]) was extracted, and mean-pooled. The original embedding is represented by

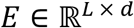

Where E is a column in the embedding, ℝ is the matrix representing the full embedding, L is the protein length and d is the dimensionality of the embedding defined by the model. The mean-pooled embedding *υ* is therefore defined by

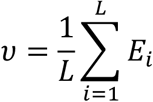

These mean pooled embeddings are then cached for later retrieval to avoid re-computation. Note that special tokens are excluded from the embedding pre-computation, such as start and stop.

#### K-means stability testing

To evaluate the stability of clustering by PLM-clust, we subsampled the CAZy families from figure 2b to take 9 subfamilies, comprising a total of 315 sequences in total. We embedded sequences using ESM2 (ESM2_t33_650M_UR50D), and ran k-means (scikit-learn, n_init=10) using a range of cluster numbers (k=2-20). For each k, clustering was repeated across 10 random seeds, and reproducibility was quantified from all pairwise replicate comparisons using adjusted Rand index (ARI) and normalized mutual information (NMI). Results were summarized using mean ± SD across all replicate pairs and visualized using Matplotlib. Controls were as follows:

1. Feature-shuffle control: each embedding dimension was independently permuted across samples, preserving per-feature marginals while disrupting multivariate structure.
2. Covariance-matched Gaussian control: synthetic embeddings were sampled from a multivariate normal with the empirical mean vector and covariance matrix of the observed embeddings.
3. Subsample stability control: 80% subsamples were drawn per replicate, k-means run on each sample, and ARI/NMI were computed on overlapping samples between replicate pairs

#### Pseudo-perplexity calculations

For one-at-a-time scoring, each sequence was tokenized and each non-special position was masked individually (excluding the start and end tokens). For each masked position, the negative log-likelihood (NLL) of the true amino acid at that site was computed from model logits, and these per-residue NLL values were averaged across the sequence to obtain one-at-a-time score.This was taken as one-at-a-time PPL*.

To calculate 15% mask score, independent masking replicates were generated using a Bernoulli mask with a probability of 0.15 at each position. Any special tokens were excluded from the masking. In each replicate, cross-entropy was computed over masked positions, and converted to PPL* as exp(loss/n_masked). The replicates scores were then averaged to obtain the 15% masking PPL* score. During benchmarking, this was run with between 1-8 replicate counts.

To calculate the single-pass score, the unmasked sequences were passed once through the model and the per-token NLL was computed directly from logits and true tokens, and averaged over all non-padding tokens to yield a single-pass PPL* score.

#### Interpretability of clustering

For interpretability analysis, 24 biophysical features were examined for their distribution within and between clusters.These metrics are outlined in Table 2.

**Table 2:**
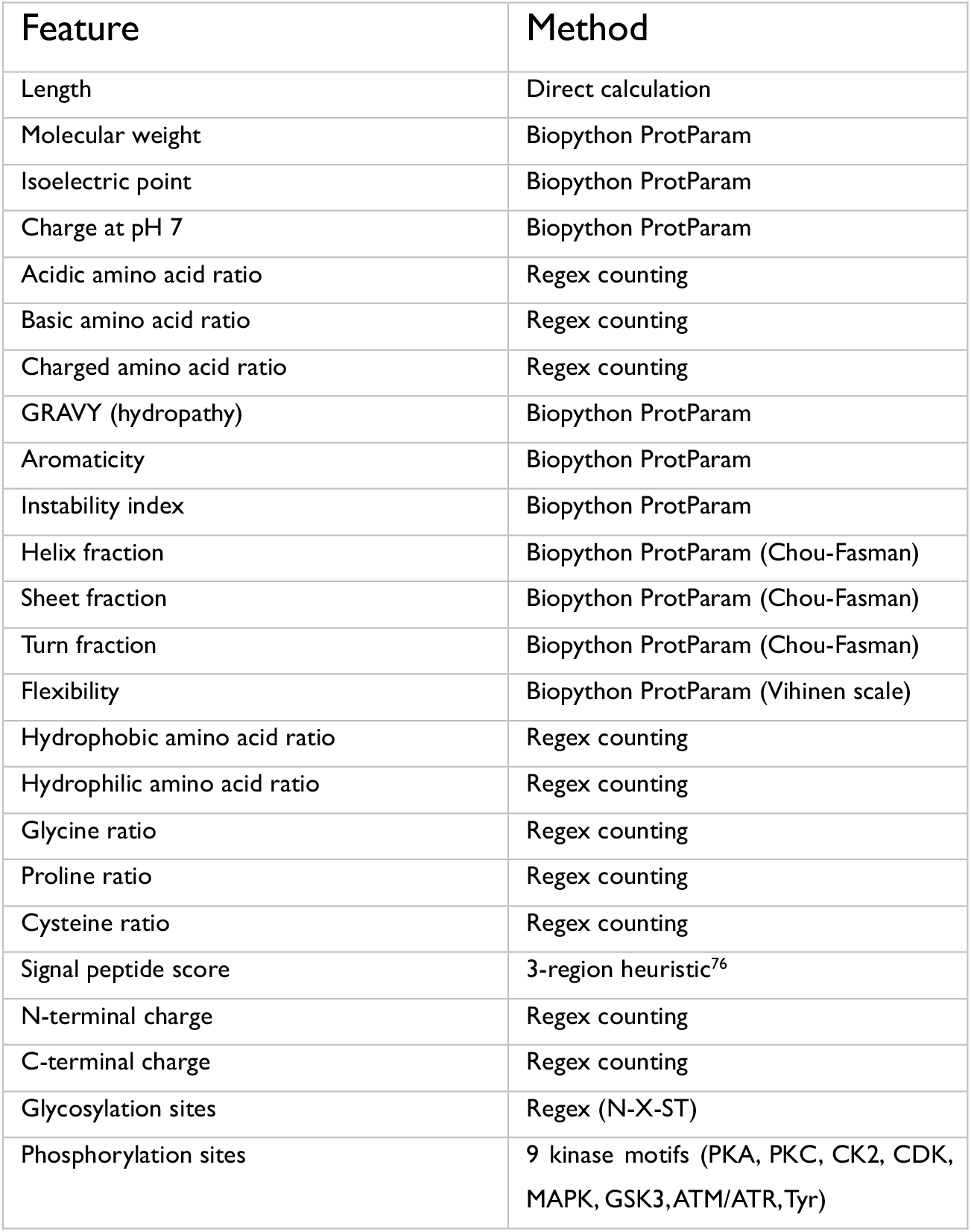
Biophysical metrics considered for interpretability of clustering.

| Feature | Method |
| --- | --- |
| Length | Direct calculation |
| Molecular weight | Biopython ProtParam |
| Isoelectric point | Biopython ProtParam |
| Charge at pH 7 | Biopython ProtParam |
| Acidic amino acid ratio | Regex counting |
| Basic amino acid ratio | Regex counting |
| Charged amino acid ratio | Regex counting |
| GRAVY (hydropathy) | Biopython ProtParam |
| Aromaticity | Biopython ProtParam |
| Instability index | Biopython ProtParam |
| Helix fraction | Biopython ProtParam (Chou-Fasman) |
| Sheet fraction | Biopython ProtParam (Chou-Fasman) |
| Turn fraction | Biopython ProtParam (Chou-Fasman) |
| Flexibility | Biopython ProtParam (Vihinen scale) |
| Hydrophobic amino acid ratio | Regex counting |
| Hydrophilic amino acid ratio | Regex counting |
| Glycine ratio | Regex counting |
| Proline ratio | Regex counting |
| Cysteine ratio | Regex counting |
| Signal peptide score | 3-region heuristic<sup>76</sup> |
| N-terminal charge | Regex counting |
| C-terminal charge | Regex counting |
| Glycosylation sites | Regex (N-X-ST) |
| Phosphorylation sites | 9 kinase motifs (PKA, PKC, CK2, CDK, MAPK, GSK3, ATM/ATR, Tyr) |

We quantified feature importance using ANOVA effect size (η^2^) for each biophysical feature across clusters. For a given feature, η^2^ was calculated as the proportion of the total variance explained by between-cluster differences^77^:

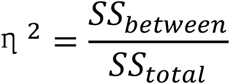

Where:

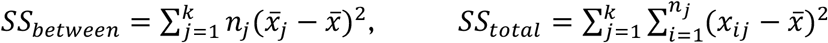

Where k is the number of clusters, n_j_ is the size of cluster j, 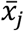 is the mean of cluster j, and 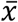 is the global mean. Note that we use η^2^ descriptively as a variance-explained measure of cluster separability, rather than as a hypothesis test.

To analyse average sequence similarity between clusters, a random set of 1000 protein pairs were taken from the dataset and subject to MAFFT alignment – average cluster identity was calculated by using the average identity within or between bins. Similarity was defined as exact-match fraction over aligned, non-gap positions, and then within/between cluster means were compared.

Phylogenetic similarity between proteins was calculated using MAFFT to create a multiple sequence alignment, followed by FastTree to generate a phylogenetic tree. Pairwise phylogenetic distance was taken as tree path length between sequence pairs.

Embedding space distance between sequence embeddings was computed from ESM2 MPEs. Cosine distance was calculated as:

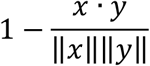

Where x and y are the pair of embeddings being compared.

#### Benchmarking variant effect prediction using ProteinGym

ProteinGym datasets were downloaded from github.com/proteingym and a random set of 100 proteins were selected from the clinical substitutions dataset (6421 total variants; 4153 pathogenic, 2269 benign). The change in the PPL* between the variant and the wild-type was calculated using single-pass scoring (above). Performance was evaluated by classifying variants as either Pathogenic or Benign, using the single-pass scoring (relative to the WT of the protein) as the predictor. ROC-AUC (scikit-learn) was computed on pooled variants across proteins.

#### Running PLM-clust for xylanases

The amino acid sequence for *Bs*GH11 was retrieved from UniProt and used as a query for NCBI’s nr dataset.The top 5000 hits by e-value were selected, and filtered by length to remove sequences with more than one domain using manual inspection of sequence lengths. The sequences were then encoded and clustered using mean-pooled embeddings and k-means,with six means, as described above. The MPEs were visualized using t-SNE (scikit-learn) and coloured by the cluster to which they belonged.The embeddings were used to calculate single-pass PPL*, as described above, and the sequence from each cluster with the lowest (i.e. best) score was suggested for experimental characterization. A suite of 24 biophysical parameters were investigated for distribution between clusters, as described above.

#### Recombinant expression and assay of xylanases

Signal peptides were identified and removed using SignalP5.0, genes codon-optimized for expression in *E. coli*, and ordered as gBlocks with 18 bp overhang of an N-terminal hexahistidine tag and C-terminal UTR of an acceptor plasmid. Genes were assembled into the plasmid using Gibson assembly and the products transformed into E. coli NEB 5a.The plasmids were extracted using silica column purification (GeneJet,Thermo), and transformed into NEB BL21 (DE3). These cells were grown at 37 °C to O.D. 0.6 in LB in 50 ml total volume, and induced with 0.5 mM IPTG.The induced cultures were shaken overnight at 20 °C. Cells were pelleted by centrifugation at 4000 xg, and the pellet resuspended in BugBuster (Millipore).The cells were shaken for 30 minutes at 20 oC.The resulting suspension was centrifuged at 12,000 xg, and the supernatant transferred to Tris-HCl (pH 8.0) with Agarose-NiNTA bead suspension (Qiagen).The mixture was left shaking for one hour, and then passed through a gravity filtration column. The matrix was washed once using Tris-HCl pH 8.0 with 5 mM imidazole, and then the protein was eluted in Tris-HCl pH 8.0 with 250 mM imidazole. Protein was concentrated and the buffer exchanged using size-exclusion filters (10 kDa, Amicon) into 100 mM Tris-HCl pH 8.0.

Endo-xylanase activity was monitored using azo-xylan (birchwood, Megazyme), by incubating purified protein (0.01 mg/ml) with 500 μl of assay mixture according to the manufacturer’s instructions. Upon 10 minutes of incubation, 500 μL of 96% EtOH was added, and the suspension centrifuged at 12,000 xg for five minutes. The supernatants were collected and measured for absorbance at 595 nm and a no-enzyme control was used to calculate baseline absorbance. Absorbance relative to the negative (no enzyme) and positive (*Bs*GH11) controls was computed and normalized by protein concentration as measured by NanoDrop microvolume spectrometer (Thermo Fisher).

Xylose release was monitored using xylose dehydrogenase coupled assay. A solution of 2 mg/ml beechwood xylan, 1 mM NAD+ and 52 U of xylose dehydrogenase/mutarotase (Megazyme) was incubated with the enzyme candidates.Absorbance was monitored at 340nm over time. The maximum rates were determined and compared to background rates, and normalized by protein concentration as measured by NanoDrop microvolume spectrometer (Thermo Fisher).These were then compared to the rate, per mg of enzyme, of *Bs*GH11.

Protein melting temperatures were determined by thermal shift assay (TSA) by incubation of 10 μg of protein with 1x Sypro Orange (Invitrogen) in Tris-HCl (pH 8.0). Sigmoidal curves were fit using the thermal shift assay fitting software from uht-biophysics (github.com/Matt115A/uht-biophysics).

#### Iterating PLM-clust for xylose-releasing enzymes

The same process was carried out as above, with some exceptions. Cluster 3 was used as the input to the clustering algorithm. Purification was carried out in a 96-well plate, rather than 50 ml cultures. Each protein was split into 8 wells of the plate. Instead of gravity columns to wash and elute the protein from the beads, centrifugation (400 xg, 3 minutes) was carried out. Assay replicates were carried out using protein from different wells of the plate. Concentration was determined by NanoDrop microvolume spectrometer (Thermo Fisher).

#### Recombinant expression and assay of imine reductases

IRED genes were codon-optimized for expression in *E. coli*, and synthesised as gBlocks (IDT) with 20 bp overhang of an N-terminal hexahistidine tag and C-terminal UTR of a custom acceptor expression plasmid (RSF origin, Extended Data 8). Genes were cloned into the plasmid using Gibson assembly and the products transformed into E. coli NEB 5a.The plasmids were extracted using silica column purification (GeneJet, Thermo). Correct assembly was verified by Sanger sequencing. The plasmids were transformed into NEB BL21 (DE3). These cells were grown at 37 °C to O.D. 0.4 in LB in 50 ml total volume and induced with 1 mM IPTG. The induced cultures were shaken overnight at 20 °C. Cells were pelleted by centrifugation at 4000 G, and the pellet resuspended in 1mL of 1x BugBuster (Millipore).The cells were shaken for 20 minutes at 20 °C.The resulting suspension was centrifuged at 16,000 xg at 4 °C and the supernatant transferred to Phosphate Buffer (50mM Phosphate, 300mM NaCl pH 8.0) with Agarose-NiNTA bead suspension (Qiagen). The mixture was left shaking for one hour and then passed through a gravity filtration column (Cytiva). The matrix was washed twice using Phosphate Buffer pH 8.0 with 30 mM imidazole, and then the protein was eluted in Phosphate Buffer pH 8.0 with 300 mM imidazole. Protein was buffer exchanged using PD-10 desalting columns (Cytiva) into Tris/HCl pH 8.0 and concentrated using size-exclusion filters (10 kDa, Amicon) into Tris-HCl/pH 8.0. Protein concentration was measured using NanoDrop microvolume spectrometer (Thermo Fisher) and by Bradford assay.

Reduction activity towards 2-methyl-1-pyrroline (2-MePy, Sigma-Aldritch) or cyclohexanone & cyclopropylamine (CHX+CPA, Sigma-Aldritch) was measured in triplicate through NADPH (Carbolution) depletion at 340 nm for 15 minutes using 0.3 mg/mL of enzyme at 21 °C. To start the reaction, 90 μL of 1.1x substrate solution was mixed with 10 μL of 10x enzyme solution to a final concentration of 10 mM ketone, 20 mM amine, and 0.5 mM NADPH in 100 mM Tris-HCl pH 8.0. Negative controls containing either no ketone or no enzyme were run in parallel. Initial rates v_o_ for NADPH depletion were obtained as the slope of time courses fitted to a linear equation and corrected for the slope of the background reaction obtained from an identical experiment without enzyme).

Protein melting temperatures were determined by thermal shift assay (TSA) by incubation of 10 μg of protein with 1x Sypro Orange (Invitrogen) in Tris-HCl (pH 8.0). Sigmoidal curves were fit using the thermal shift assay fitting software from uht-biophysics (github.com/Matt115A/uht-biophysics).

To iterate on the process, the cluster containing the best performing 2-MePy reductase was further subclustered and 5 more cluster representatives were characterised as above. Non-natural sequences were removed and the second-best hit tested, if they constituted a cluster representative.

#### Generalization between models

Sequence embeddings were extracted for GH11 proteins. Using five protein language models: ESM2 (8M, 35M, 150M and 650M parameter variants) and ProtBERT^57^. For each model, sequence-wise embeddings were obtained using mean-pooling and clustered using k-means (k=6). Between-model agreement was quantified with ARI and NMI, between model-specific cluster assignments.

As a control, we used a feature-shuffle embedding null: starting from the ESM2 650M embedding matrix, each embedding dimension was independently permuted across sequences (preserving per-dimension marginal distributions while disrupting multivariate structure), followed by k-means clustering (k=6).

For the cross-model visualization panel, 100 GH11 sequences were sampled from the GH11 campaign. t-SNE was computed on ESM-2 650M embeddings, and points were then coloured by either their ESM-2 or ProtBERT k-means labels.

## Supporting information

Supplementary Information Penner et al

## Data availability

All raw data and processing scripts used to generate each figure in this manuscript are available at github.com/Matt115A/plmclust-benchmarking-public.git.The codes can be run locally using make *plm-clust-manuscript* to re-generate plots as presented here.

## Author Contributions

M.P. designed and ran the algorithm and benchmarking criteria, and carried out molecular biology, expression and enzyme screening of GH11 enzymes. Melting temperature measurements were carried out by H.M and P.N. M.L and H.B. carried out molecular biology, expression and enzyme screening of IREDs. M.L. ran PLM-clust for the IREDs. P.D. and F.H. conceptualized the project and provided experimental design and writing.

## Acknowledgements

The authors would like to thank Simon Mathis and Kieran Didi for their valuable discussions.

## Supplementary Information

A. Supplementary Figures 1-12
  1. SSN of the CAZymes used in this study alongside general principle of SSN generation
  2. Comparison of embedding distance metrics with phylogenetic distance
  3. Comparing MPEs and SWEs for GH1
  4. Melting curves of GH11 enzymes from iteration 1
  5. Partial sequences annotated per cluster in iteration 1
  6. Melting curves of the top hit from the GH11 campaign
  7. Time-calibrated phylogeny of *Bacillales* members in the GH11 campaign
  8. Sequence identity of screened IREDs in iteration 1
  9. Melting curves of IRED enzymes from iteration 2
  10. Single-pass PPL* correlations with one-at-a-time masking on diverse CAZymes for different PLMs
  11. Google Colab interface for PLM-clust
  12. Web interface for PLM-clust
B. Supplementary Tables 1-3
  1. GH families with best and worst agreement between phylogeny and MPE
  2. Features and their effect sizes for GH11
  3. Features and their effect sizes for *Sr*IRED

## Extended Data

*All extended data are provided as uploaded .zip files*

Extended data 1: CAZy sequences used in this study, with their calculated cluster identities and PPL* scores

Extended data 2: GH1 sequences used in this study, with their calculated cluster identities and PPL* scores

Extended data 3: GH sequences used in this study for phylogenetic comparisons, with their calculated cluster identities and PPL* scores

Extended data 4: GH11 enzymes used in this study in round 1, with their calculated cluster identities and PPL* scores

Extended data 5: GH11 enzymes used in this study in round 2, with their calculated cluster identities and PPL* scores

Extended data 6: Correlation coefficients between phylogeny and cosine similarity for selected CAZy families

Extended data 7: GH11 enzymes tested in this study

Extended data 8: IRED enzymes used in this study, with their associated plasmid maps Extended data 9: Sequence Similarity network of the selected members of the CAZY database Extended data 10: Parameters used to generate the sequence similarity network

