## Supplementary Information Penner et al for "Unsupervised protein language models learn patterns of enzyme function": Penner_23April_SI.pdf

**Authors:** Matthew Penner[1], Michal Lihan[1], Hannes Bormke[1], Peter Nix[1], Hanna Moscho[1], Paul Dupree[1,*] and Florian Hollfelder[1,*]

**Affiliations:** [1]Department of Biochemistry, University of Cambridge, 80 Tennis Court Road, Cambridge, CB2 1GA, UK

---

Contents:

A. Supplementary Figures 1-12
   1. SSN of the CAZymes used in this study alongside general principle of SSN generation
   2. Comparison of embedding distance metrics with phylogenetic distance
   3. Comparing MPEs and SWEs for GH1
   4. Melting curves of GH11 enzymes from iteration 1
   5. Partial sequences annotated per cluster in iteration 1
   6. Melting curves of the top hit from the GH11 campaign
   7. Time-calibrated phylogeny of *Bacillales* members in the GH11 campaign
   8. Sequence identity of screened IREDs in iteration 1
   9. Melting curves of IRED enzymes from iteration 2
   10. Single-pass PPL* correlations with one-at-a-time masking on diverse CAZymes for different PLMs
   11. Google Colab interface for PLM-clust
   12. Web interface for PLM-clust

B. Supplementary Tables 1-3

C. Extended data descriptions 1-10

### Supplementary Figures

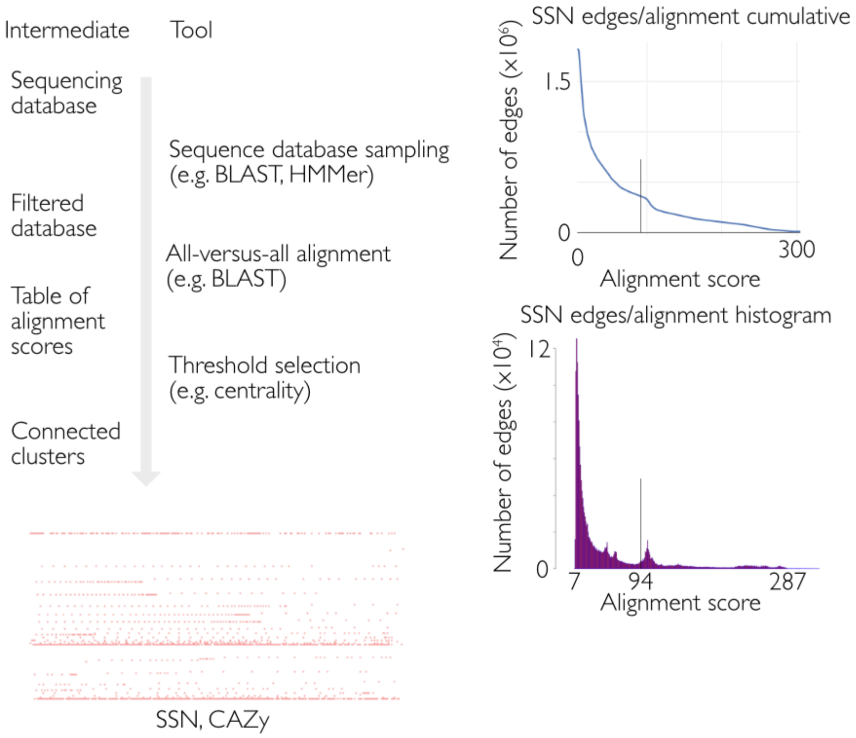


Figure S1: SSN of the CAZymes used in this study alongside general principle of SSN generation. (A) Pipeline for generation of an SSN (B) cumulative and histogram plots of edge number compared to alignment score for the CAZy subset used in this work (C) Visualization of the network thus derived using a force-directed layout in Cytoscape.

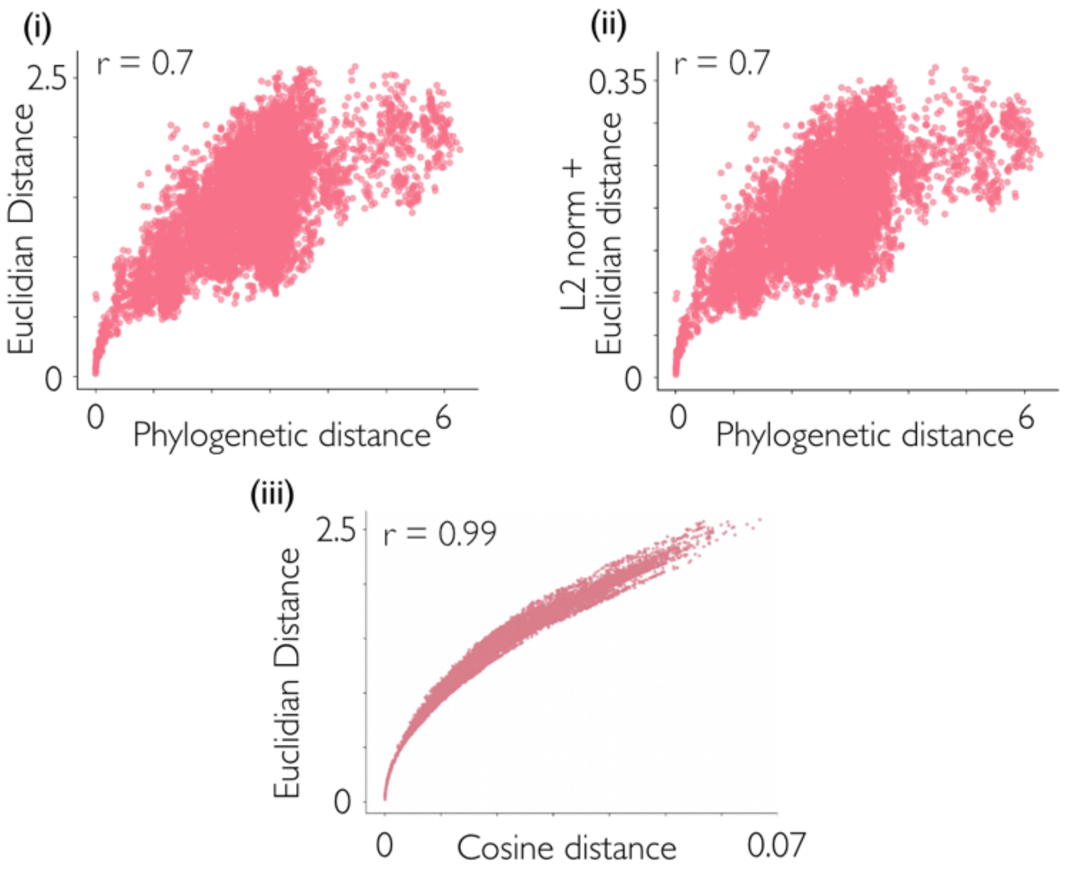


Figure S2: (i) Relationship of Euclidian distance of GH1 MPEs with phylogenetic distance. N=10,000 sequence pairs (Spearman r = 0.70)(ii) Relationship of L2 regularization + Euclidian distance of GH1 MPEs with phylogenetic distance, with the same sequence pairs (Spearman r = 0.69). (iii) Relationship of the same distances as measured using cosine/Euclidian distance (without L2 regularization, Spearman r = 0.99)

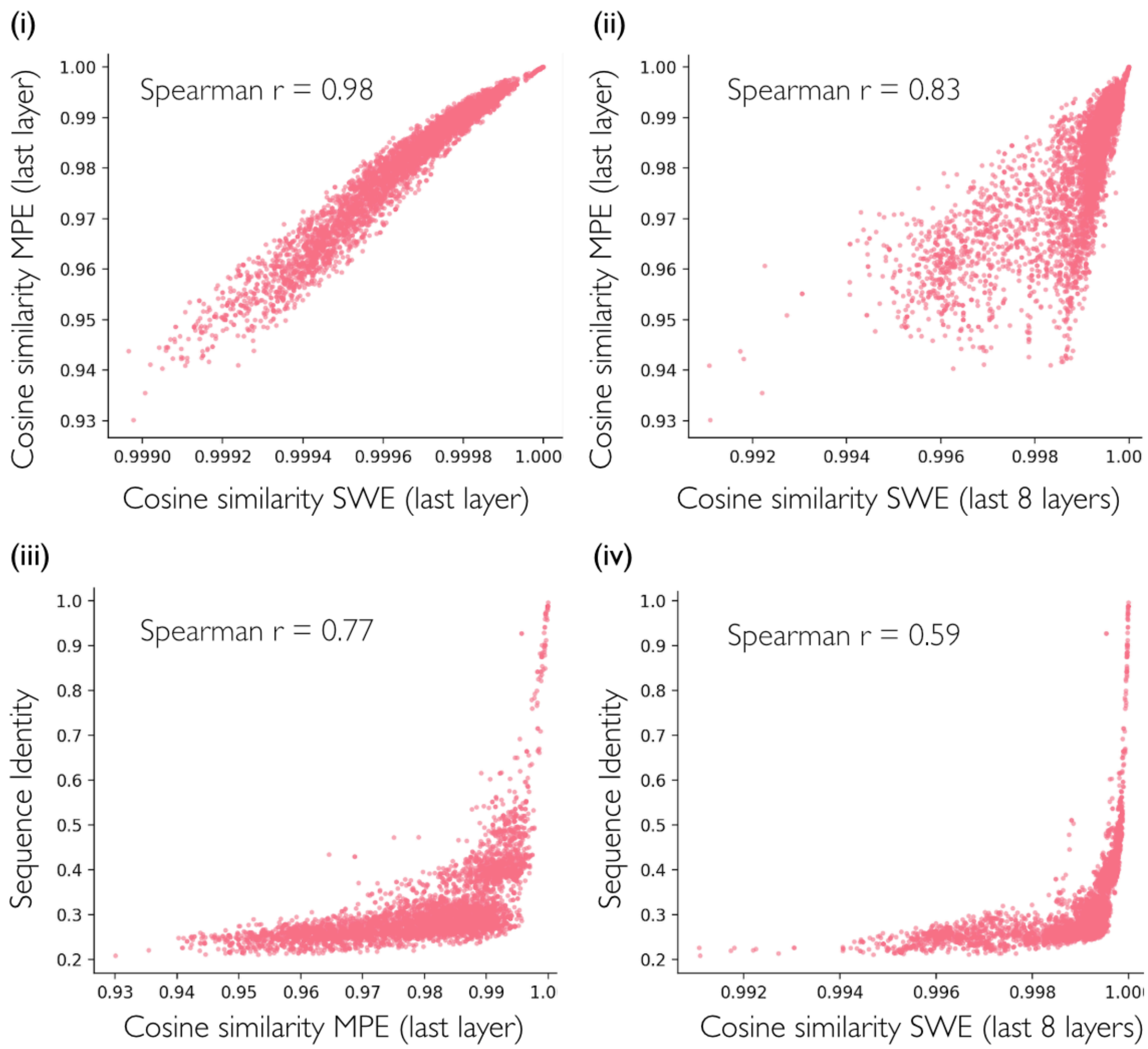


Figure S3: (i) Relationship between cosine similarity of a random set of 10,000 pairs of GH1 enzymes computed by MPEs on the last layer of ESM-650 embeddings (as used in PLM-clust), and cosine similarities of the same pairs by SWEs on the final layer of the embeddings. Spearman r = 0.98. (ii) As (i), but SWEs using the final 8 layers of the embedding. Spearman r = 0.83. (iii) Relationship between sequence identity and MPE similarity of the same pairs (Spearman r = 0.77) and (iv) as (iii) but using SWE similarity on the final 8 embedding layers (Spearman r = 0.59).

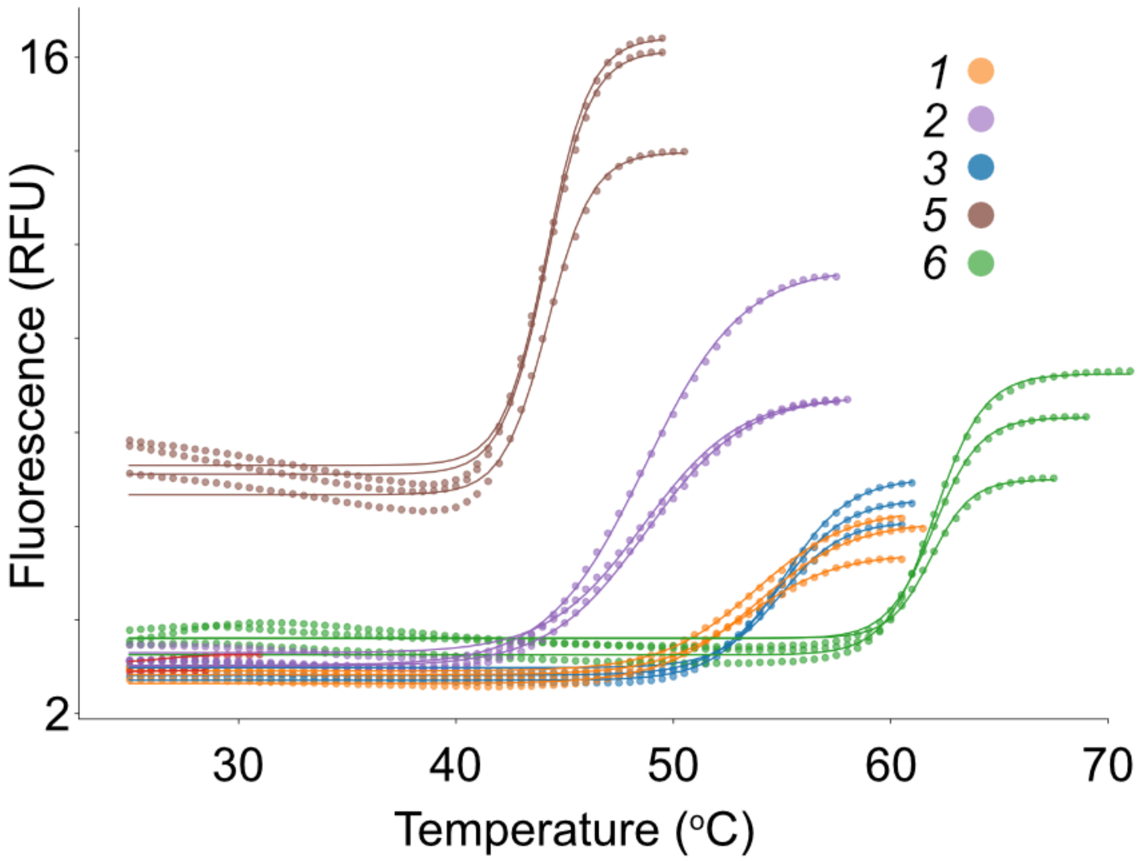


Figure S4: Melting curves and sigmoid fits of GH11 round 1 enzymes used in this study, derived by thermal shift assay (TSA). Curves were fit using the uht-biophysics pipeline (methods), and error in the $T_m$ was estimated using the standard deviation of the three inflection points for each mutant.

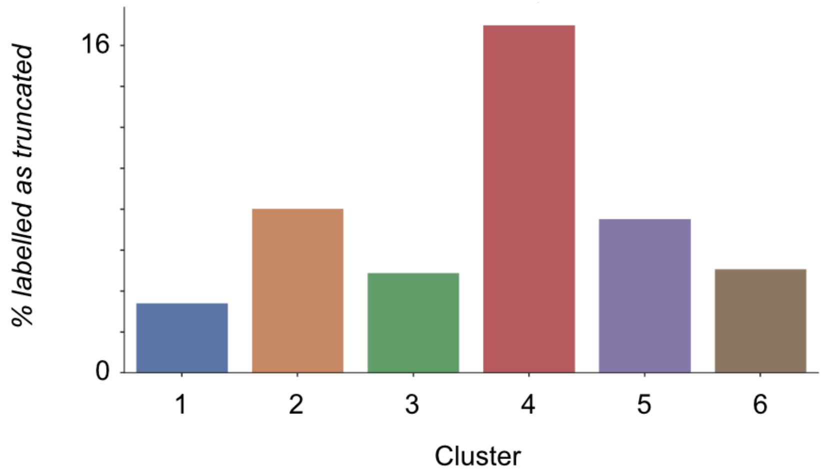


Figure S5: GH11 clusters and their proportion of annotated 'partial' in their NCBI sequence headings. These were identified using Regex for the phrase 'partial' in the headers of the sequences downloaded from NCBI. Percentage represents the percentage of sequences in each of the clusters.

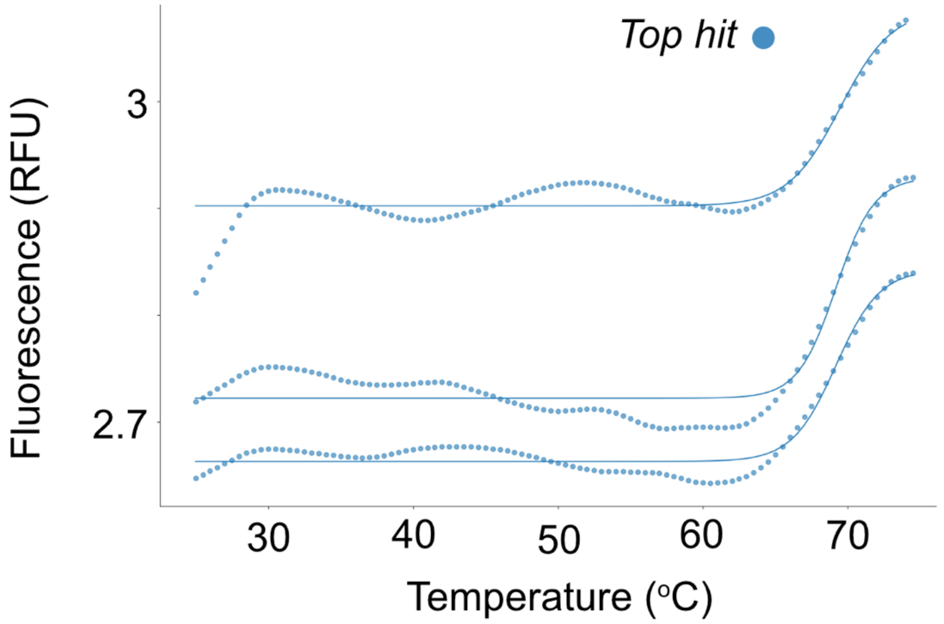


Figure S6: Tm curves of the top GH11 hit found in this study, derived by thermal shift assay (TSA). Curves were fit using the uht-biophysics pipeline (methods), and error in the $T_m$ was estimated using the standard deviation of the three inflection points for the enzyme.

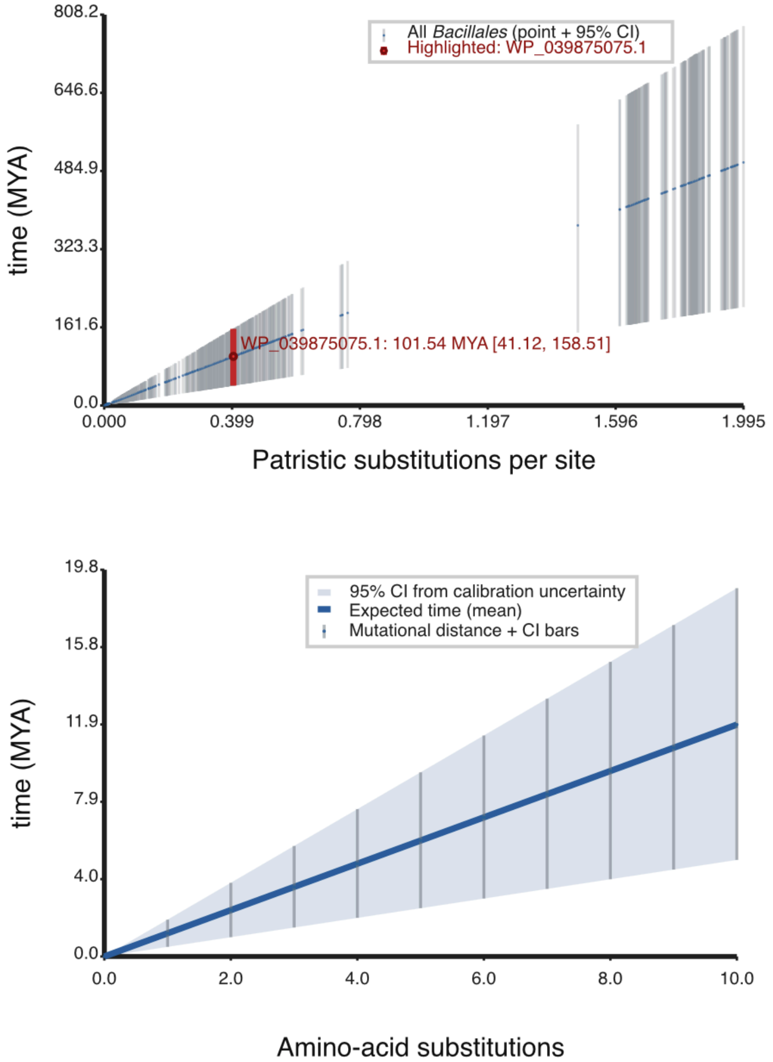


Figure S7: Time-calibrated phylogeny of *Bacillales* members in the GH11 campaign, in distance from BsGH11. (i) WP_039875075.1, the cluster representative for the cluster that BsGH11 is found in, is highlighted in red. (ii) Using the calibration to estimate evolutionary time spanned by a low number of substitutions in the background of *Bs*GH11. Dcal = 1.995 substitutions per site, Tcal = 503 ± 150 MYA assumed calibration time between BsGH11 and the calibration marker. MYA: millions of years ago.

**Percent Identity Matrix**

| | | | | | | | |
|---|---|---|---|---|---|---|---|
| cluster=1 | 100.00% | 43.21% | 48.24% | 48.94% | 50.87% | 48.43% | 51.57% |
| cluster=4 | 43.21% | 100.00% | 48.56% | 56.83% | 51.76% | 50.18% | 50.00% |
| cluster=3 | 48.24% | 48.56% | 100.00% | 56.49% | 57.24% | 53.68% | 57.69% |
| cluster=0 | 48.94% | 56.83% | 56.49% | 100.00% | 59.86% | 58.45% | 58.25% |
| cluster=5 | 50.87% | 51.76% | 57.24% | 59.86% | 100.00% | 54.48% | 62.41% |
| cluster=2 | 48.43% | 50.18% | 53.68% | 58.45% | 54.48% | 100.00% | 69.90% |
| srIRED | 51.57% | 50.00% | 57.69% | 58.25% | 62.41% | 69.90% | 100.00% |

Figure S8: Sequence identity of screened IREDs in iteration 1. Sequence identities calculated and visualized using Uniprot web tool.

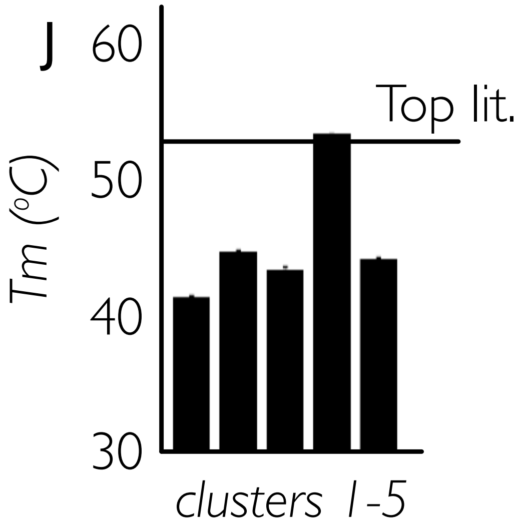


Figure S9: $T_m$ of IREDs from iteration 2. Error bars represent one standard deviation of independent replicates.

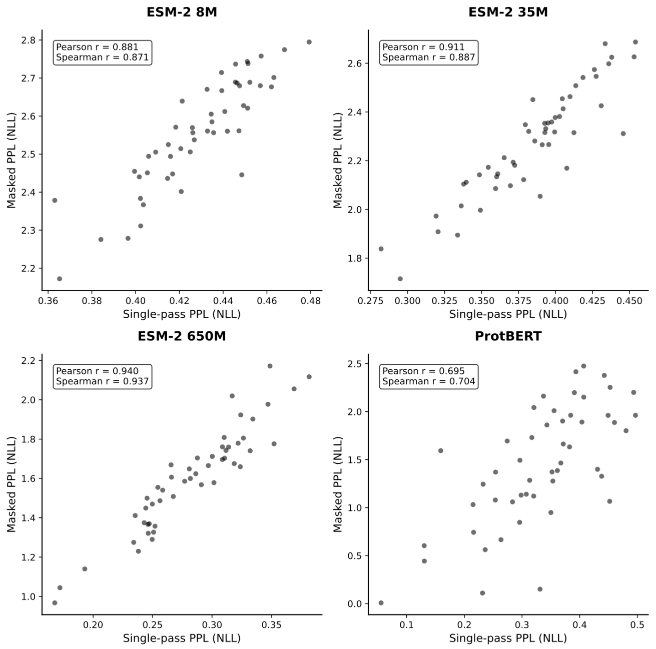


Figure S10: Single-pass PPL* correlations with one-at-a-time masking on diverse CAZymes for different PLMs. Spearman and Pearson r are reported in each case.

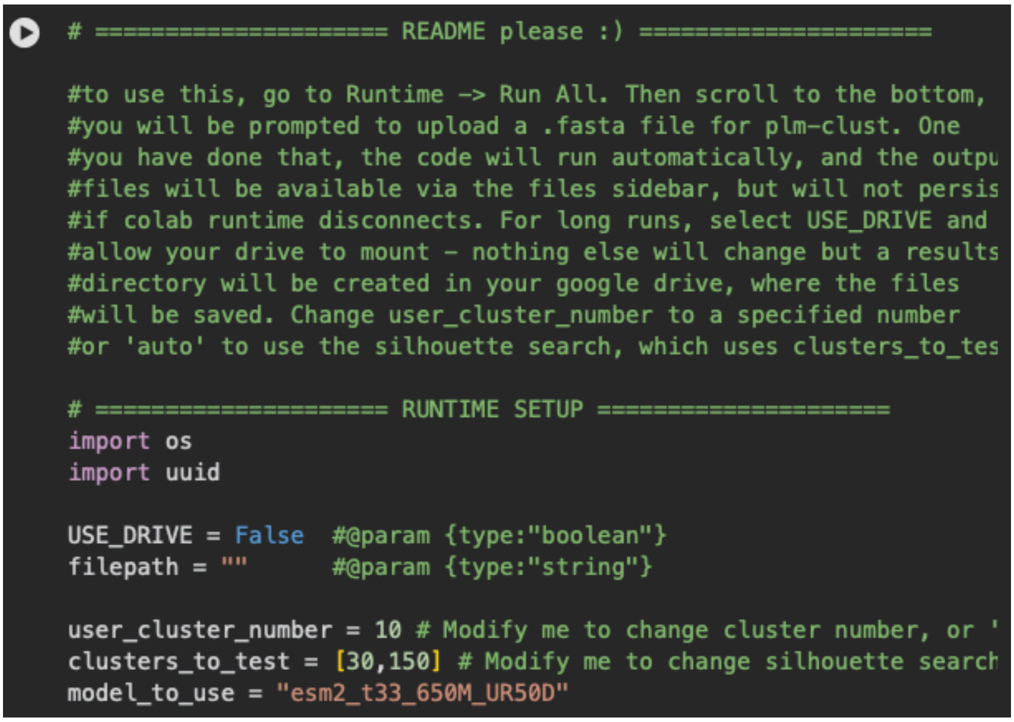


Figure S11: Google Colab interface. Users can use Google colab versions to run PLM-clust on their own data, using compute provided by the service.

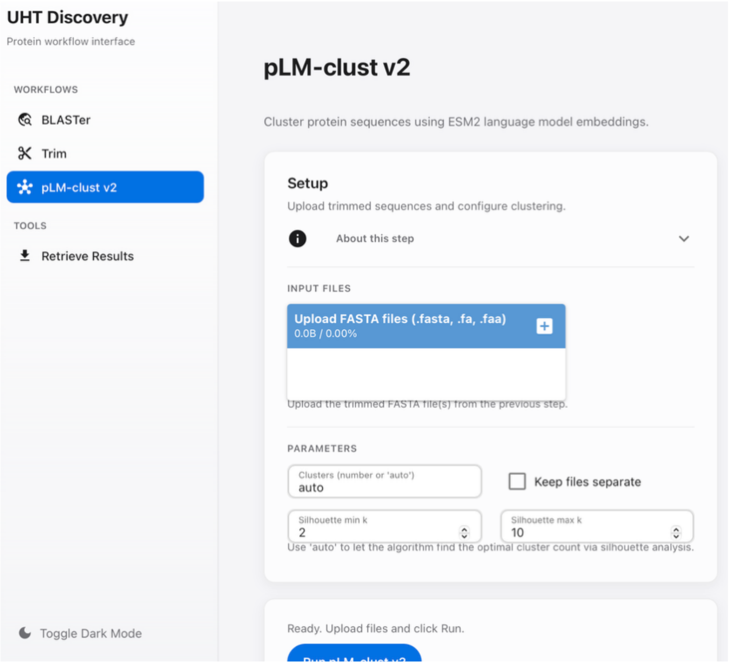


Figure S12: Web interface. The web interface can be used for all three stages – sequence retrieval, quality control and PLM-clust. It also allows for data exploration of the results via interactive t-SNE and UMAP representations of the final embeddings.

---

### Supplementary Tables

| GH family (best agreement) | Spearman r MPE vs phlogeny | Sequences sampled | | GH family (worst agreement) | Spearman r MPE vs phylogeny | Sequences sampled |
|---|---|---|---|---|---|---|
| GH193 | 0.96 | 11 | | GH31 | 0.05 | 51 |
| GH137 | 0.95 | 57 | | GH16_1 | 0.09 | 157 |
| GH2_8 | 0.95 | 80 | | GH115 | 0.089 | 146 |
| GH17 | 0.94 | 50 | | GH7 | 0.11 | 152 |

Table S1: GH families with best and worst agreement between phylogeny and MPE distance

| Feature Name | Effect size (eta squared) |
|---|---|
| N-glycosylation sites | 0.66 |
| Protein isoelectric point | 0.62 |
| Protein charge at pH 7 | 0.60 |
| Acidic amino acid ratio | 0.51 |

| | |
|---|---|
| Basic amino acid ratio | 0.43 |
| Glycine ratio | 0.38 |
| Charged amino acid ratio | 0.37 |
| Alpha helix fraction | 0.37 |
| Signal peptide probability | 0.34 |
| Protein aromaticity | 0.28 |
| Protein flexibility | 0.27 |
| Beta turn fraction | 0.26 |
| N-terminal charge | 0.26 |
| Beta sheet fraction | 0.22 |
| Protein sequence length | 0.22 |
| Protein molecular weight | 0.21 |
| Proline ratio | 0.21 |
| Hydrophobic amino acid ratio | 0.21 |
| Grand Average of Hydropathy | 0.18 |
| Predicted phosphorylation sites | 0.18 |
| C-terminal charge | 0.15 |
| Protein instability index | 0.15 |
| Cysteine ratio | 0.12 |
| Hydrophilic amino acid ratio | 0.09 |

Table S2: Features and their effect sizes for GH11

| Feature Name | Effect size (eta squared) |
|---|---|
| Proline ratio | 0.45892675 |
| C-terminal charge | 0.37342111 |
| Hydrophilic amino acid ratio | 0.35427419 |
| Protein isoelectric point | 0.34204622 |
| Protein aromaticity | 0.32897964 |
| Protein charge at pH 7 | 0.2893303 |
| Beta turn fraction | 0.27479945 |
| Glycine ratio | 0.27100677 |
| Acidic amino acid ratio | 0.25117253 |
| Protein flexibility | 0.25105094 |
| Alpha helix fraction | 0.24674029 |
| Beta sheet fraction | 0.20212129 |
| Predicted phosphorylation sites | 0.1706534 |
| Hydrophobic amino acid ratio | 0.16127405 |
| Protein instability index | 0.16042295 |

| | |
|---|---|
| Basic amino acid ratio | 0.15509749 |
| Grand Average of Hydropathy | 0.15429071 |
| N-glycosylation sites | 0.14918551 |
| Protein molecular weight | 0.07534425 |
| Protein sequence length | 0.05476289 |
| Signal peptide probability | 0.04510832 |
| Charged amino acid ratio | 0.02715196 |
| N-terminal charge | 0.01996422 |
| Cysteine ratio | 0.01904541 |

Table S3: Features and their effect sizes for *Sr*IRED

---

### Extended data

*All extended data are provided as uploaded .zip files*

Extended data 1: CAZy sequences used in this study, with their calculated cluster identities and PPL* scores

Extended data 2: GH1 sequences used in this study, with their calculated cluster identities and PPL* scores

Extended data 3: GH sequences used in this study for phylogenetic comparisons, with their calculated cluster identities and PPL* scores

Extended data 4: GH11 enzymes used in this study in round 1, with their calculated cluster identities and PPL* scores

Extended data 5: GH11 enzymes used in this study in round 2, with their calculated cluster identities and PPL* scores

Extended data 6: Correlation coefficients between phylogeny and cosine similarity for selected CAZy families

Extended data 7: GH11 enzymes tested in this study

Extended data 8: IRED enzymes used in this study, with their associated plasmid maps

Extended data 9: Sequence Similarity network of the selected members of the CAZY database

Extended data 10: Parameters used to generate the sequence similarity network
